# Evidence Accumulation in Synaptic Efficacy: Activity-Silent Decision-Making with Short-Term Plasticity

**DOI:** 10.64898/2026.09.27.754756

**Authors:** Tianhao Chu, Yuling Wu, Chenhao Zhang, Fang Fang, Si Wu

**Author notes:** These authors contributed equally to this work (Tianhao Chu, Yuling Wu).

## Abstract

Decision-making requires accumulation of evidence over time, yet exactly how neural circuits implement this computation remains debated. Classical models posit that evidence accumulation is realized by ramping neural activity, whereas recent experimental findings suggest an alternative mechanism in which neural activity remains largely silent before abruptly transitioning into an active decision state. This mechanism leaves a fundamental question unanswered: how is evidence accumulated during the neural activity-silent period? To address this question, we trained excitatory–inhibitory recurrent neural networks (RNNs) with short-term synaptic plasticity (STP) on a reaction-time decision task. We found that after training, the networks learned to leverage STP to make decisions in an activity-silent manner. Disrupting STP impaired decision performance, and decoding revealed a trace of accumulated sensory evidence in synaptic efficacy. Choice-tuning analysis revealed a clear functional organization of the network, and connection-pruning analysis uncovered the coupling between choice-selective excitatory and inhibitory neural groups. Guided by the architecture of trained RNNs, we constructed a minimal neural circuit model, in which evidence accumulates in facilitated synaptic efficacy that progressively reshapes the circuit’s phase portrait, eventually driving the network state across the stability boundary and triggering a rapid transition to a choice state. We further compared activity before a choice between the theoretical STP and Static models to examine the potential energy benefit of activity-silent decision-making. Together, our results suggest STP as a mechanism for evidence accumulation in activity-silent decision-making and offer insights into the design of energy-efficient artificial neural networks.

## 1 Introduction

Decision-making often requires accumulating sensory evidence before making a choice. Early experimental and theoretical studies suggested that neurons representing competing choices accumulate evidence through a gradual increase in firing rate, known as ramping [Roitman and Shadlen, 2002, Wang, 2002, Wong and Wang, 2006]. However, more recent studies have identified a different activity pattern, known as stepping, in which firing changes abruptly during a decision [Latimer et al., 2015, Zoltowski et al., 2019]. The activity of neurons representing the eventual choice remains low after stimulus onset and then rises rapidly before the choice is reported. This pattern resembles activity-silent working memory, in which information is retained with little sustained neuronal activity [Rose et al., 2016, Wolff et al., 2017]. We refer to the low-activity interval followed by a rapid choice-related rise as *activity-silent decision dynamics*. Ramping-based accounts link evidence accumulation to rising neuronal activity, so activity-silent decision dynamics raise two questions: how does the circuit accumulate evidence while choice-related activity remains low, and what dynamics keep neuronal activity low after stimulus onset and then cause it to rise rapidly?

Training artificial neural networks on behavioral tasks has become an established approach to studying neural computation [Mante et al., 2013, Song et al., 2016]: optimizing a task objective can produce an activity pattern without prescribing the internal activity trajectories [Cueva and Wei, 2018, Banino et al., 2018], and analysis of the trained network then identifies the neurons and connections that generate the pattern and support behavior [Roach et al., 2023, Chu et al., 2026]. We use this approach to investigate whether task optimization produces activity-silent decision dynamics and how the resulting circuit performs evidence accumulation.

We hypothesize that short-term synaptic plasticity (STP) supports evidence accumulation during activity-silent decision dynamics. STP is a widespread property of biological synapses in which recent presynaptic activity transiently increases or decreases synaptic efficacy, often through changes in neuro-transmitter release [Zucker and Regehr, 2002, Regehr, 2012]. Previous models have shown that STP can support working memory and stimulus discrimination [Mongillo et al., 2008, Masse et al., 2019, Bourjaily and Miller, 2012, Ballintyn et al., 2019]. Here, we test whether a recurrent neural network (RNN) trained for decision-making learns to recruit STP while choice-related neuronal activity remains low. We introduce STP into an excitatory–inhibitory (E–I) RNN with separate excitatory (E) and inhibitory (I) neurons and train it on a motion-discrimination task [Song et al., 2016]. Each presynaptic neuron has learnable STP parameters that are optimized during training and held fixed within each trial, while synaptic efficacy evolves throughout the trial to modulate recurrent transmission. We compare RNNs with learnable STP against Static networks trained without STP.

We found that RNNs with learnable STP consistently developed activity-silent decision dynamics after training across initialization conditions. These dynamics were more prevalent than in Static networks and were accompanied by greater recruitment of STP. The transition time, defined as the onset of the rapid rise in neuronal activity, varies across trials, so averaging neuronal activity across trials produces ramping, consistent with experimental observations [Latimer et al., 2015]. Perturbing STP impairs decisions with learned weights held fixed, and synaptic variables provide a more accessible linear readout of accumulated sensory evidence than neuronal activity. Neurons with higher STP strength are predominantly choice-selective, and silencing them impairs performance more than silencing an equal number of low-STP neurons with matched E/I composition. Connection pruning identifies a clear choice-selective E–I circuit. Excitatory neurons reinforce the same choice and recruit inhibitory neurons with the same choice preference; those inhibitory neurons suppress excitatory neurons with the opposite preference. These results locate strong STP recruitment within the circuit that supports the decision.

Although the trained RNNs with STP developed activity-silent decision-making dynamics similar to those seen in neural recordings, the dynamical mechanism that produces these patterns in the trained networks remains largely opaque. Guided by the choice-selective circuit organization identified in the RNNs, we therefore developed a theoretical model to explain how choice-related firing can remain low and then rise abruptly, as observed experimentally. We incorporate this organization and population-level STP into a classical competitive decision framework [Wang, 2002, Wong and Wang, 2006]. The model has two competing choice groups, each with excitatory and inhibitory populations that support self-excitation and suppression of the opposing group. As synaptic efficacy increases, the corresponding Static network passes from a tristable regime, in which a stable low-activity state coexists with two stable choice states, to a bistable regime in which the low-activity state becomes a saddle point [Wong and Wang, 2006]. In the STP model, synaptic efficacy evolves over time: its initially low value keeps neuronal activity low after stimulus onset while evidence accumulates in synaptic efficacy. Once efficacy crosses a critical value, the low-activity state loses stability and activity in the winning group rises rapidly, explaining both how evidence accumulates during low neuronal activity and why it ends in a rapid transition. Comparison with a Static ramping model illustrates the potential energy benefit of keeping activity low before a choice.

Our contributions are threefold. We establish a task-trained neural network model that exhibits activity-silent decision dynamics with STP supporting evidence accumulation and identify its choice-selective E–I circuit. We propose a theoretical model in which evidence accumulates in synaptic efficacy and STP drives a transition from tristability to bistability by destabilizing the low-activity state. We connect this account to an activity-cost comparison relevant to the design of energy-efficient AI decision systems.

## 2 Task design and model architecture

We trained E–I RNNs on the reaction-time motion-discrimination task of Song et al. [2016] (Figure 1b). Two inputs provide noisy evidence for leftward and rightward motion, and a third input signals stimulus presentation. The network reports its choice through one of two outputs. Higher motion coherence increases the difference between the mean evidence inputs, making the direction easier to identify. Networks are trained to minimize the mean squared error between the two outputs and piecewise-constant choice targets during the supervised response interval (Equation 9). Appendix A.1 details the task and behavioral measures.

**Figure 1:**
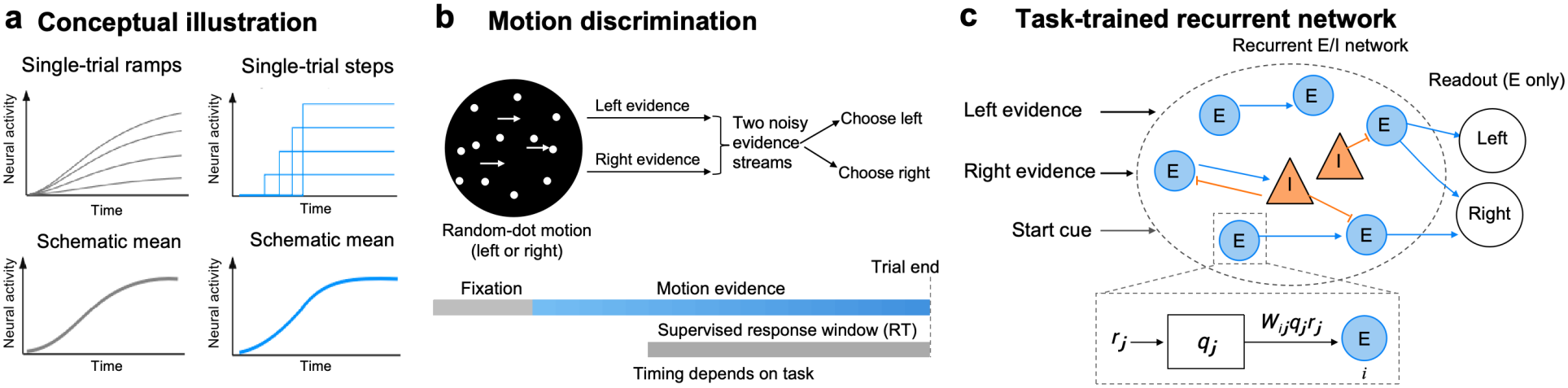
Task and network overview. **a**, Schematic ramping and activity-silent dynamics. Variation in transition times across trials can produce a gradual trial average. **b**, Reaction-time motion discrimination. The network receives noisy leftward and rightward motion evidence and reports a choice through one of two outputs. Motion coherence controls the difference between the mean evidence inputs. **c**, E–I RNN with learnable STP. STP modulates recurrent transmission, and the two choice outputs receive input from excitatory neurons. Connectivity is schematic. Appendices A.1 and A.2 specify the task and network implementation.

Each RNN contains 80 excitatory neurons and 20 inhibitory neurons, represented by nonnegative rate units. Excitatory neurons make nonnegative recurrent connections, and inhibitory neurons make nonpositive recurrent connections. Two linear choice outputs receive input from the excitatory neurons (Figure 1c). Neuronal activity follows

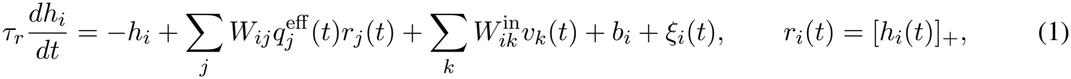

where *h*_*i*_ is the state of neuron *i, r*_*i*_ is its activity in arbitrary units, and [*h*_*i*_]_+_ = max(0, *h*_*i*_) is the rectified-linear activation function. The parameter *τ*_*r*_ is the neuronal time constant. The weight *W*_*ij*_ connects presynaptic neuron *j* to neuron *i*, 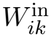 connects external input *v*_*k*_ to neuron *i*, and *b*_*i*_ and *ξ*_*i*_ are the bias and recurrent noise. The factor 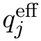 scales all recurrent outputs from neuron *j*.

STP changes recurrent transmission through two variables for each presynaptic neuron: synaptic utilization *u*_*j*_ and the available-resource fraction *x*_*j*_. They evolve according to

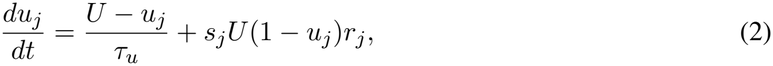

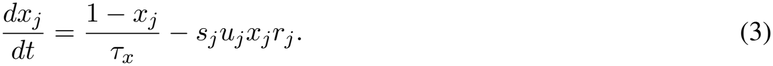

Here, *U* is baseline synaptic utilization, and *τ*_*u*_ and *τ*_*x*_ are the recovery time constants for *u*_*j*_ and *x*_*j*_. The learnable rate scale *s*_*j*_ converts neuronal activity into the rate driving STP dynamics. The synaptic efficacy normalized by its resting value is *u*_*j*_*x*_*j*_*/U*.

We compare two conditions. Static networks use 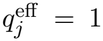 . In RNNs with STP, the learnable STP strength *a*_*j*_ determines how strongly normalized synaptic efficacy changes recurrent transmission:

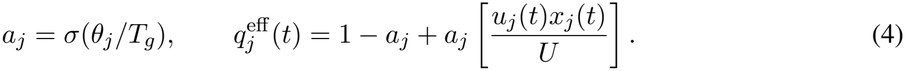

The logistic function *σ* maps the trainable parameter *θ*_*j*_ to an STP strength between zero and one. The parameter *T*_*g*_ sets the scale of this mapping. The limits *a*_*j*_ = 0 and *a*_*j*_ = 1 correspond to static transmission and full normalized synaptic efficacy, respectively. Training adjusts *a*_*j*_ and *s*_*j*_ together with the network weights, while *u*_*j*_ and *x*_*j*_ evolve within each trial. Both conditions use the same neuronal dynamics, task, and readout. Appendix A.2 gives parameter values, initialization, numerical integration, and equivalence checks. Appendix A.3 describes training and checkpoint selection, and Appendix A.4 summarizes the paired experimental configuration.

## 3 Activity-silent decision dynamics emerge in RNNs with STP

We found that activity-silent decision dynamics emerged more frequently in RNNs with STP. Training reduced task loss and improved task performance (Figure 2a). After training, among choice-tuned neurons, activity-silent dynamics were more prevalent in RNNs with STP (52.2 ± 16.2%) than in Static RNNs (36.2 ± 7.0%; mean ± standard deviation (SD) across networks), shifting the population toward the high prevalence of step-like single-trial dynamics reported in choice-selective LIP neurons (78%; Latimer et al., 2015). Selected single-trial examples further illustrate this contrast, showing ramping versus activity-silent responses under identical sensory input and noise (Figure 2b; Appendix A.5).

**Figure 2:**
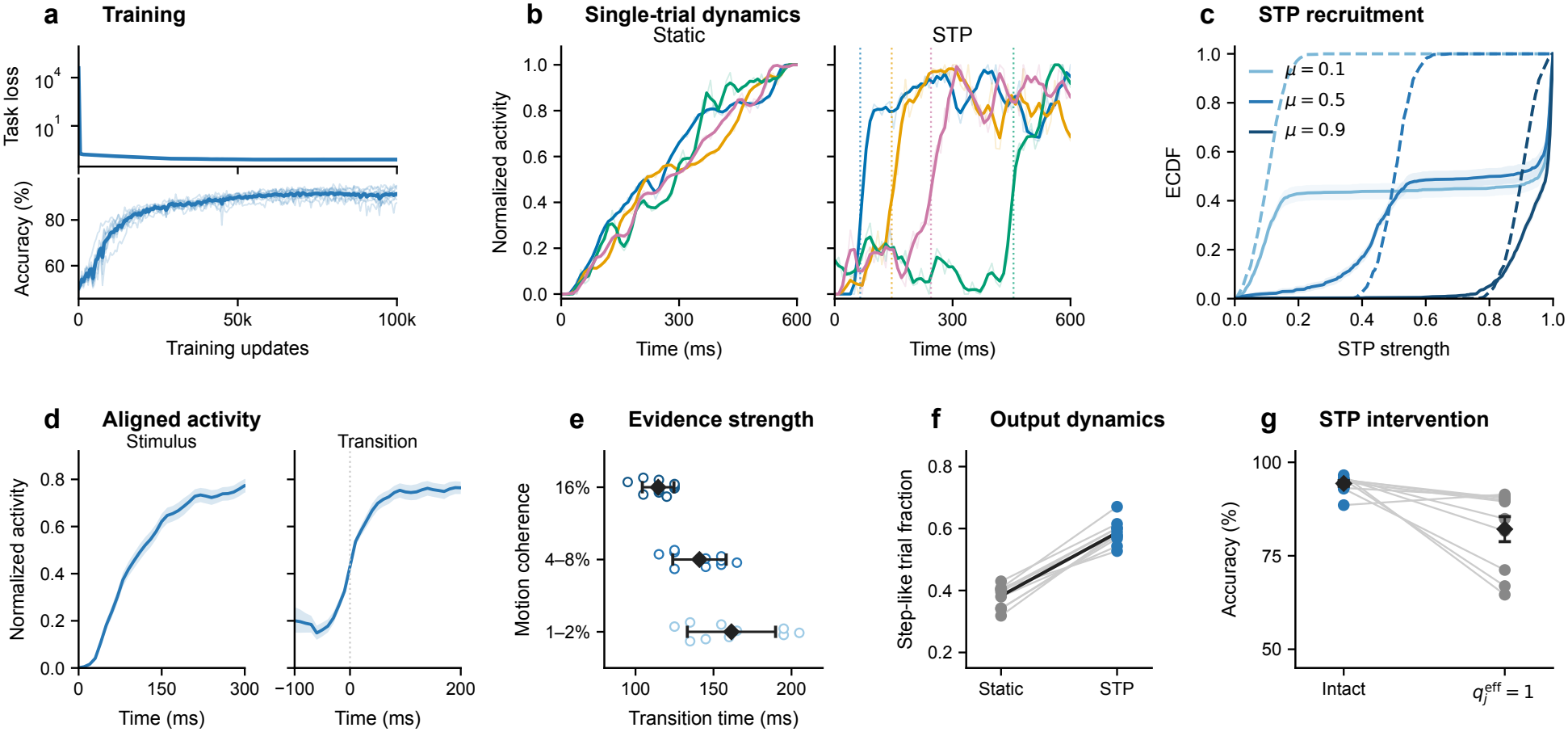
Learning and activity-silent decision dynamics. **a**, Validation loss and performance during training for the ten RNNs with STP. **b**, Matched single-trial examples showing ramping in the Static RNN and activity-silent decision dynamics in the RNN with STP under the same sensory input and noise. **c**, STP-strength distributions before (dashed) and after training (solid). Training increases STP strength in subsets of neurons under each initialization condition. **d**, The same trials produce a gradual ramp when aligned to stimulus onset and a rapid rise when aligned to their fitted transition times. **e**, Transitions occur later under weaker sensory evidence (within-network median transition times of representative units). **f**, Fraction of choice-output trajectories classified as step-like in Static networks and RNNs with STP. **g**, Final-choice accuracy in the intact STP network and with effective synaptic efficacy set to one 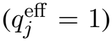 , while trained weights are held fixed. Lines join paired networks; diamonds and error bars show mean ± SEM across ten networks. Panels a, c, and e–g summarize ten networks (per initialization in c); b and d show single-network examples. Analysis details are given in Appendices A.5 and A.7.

We next asked whether task optimization alone leads networks to exploit STP. Comparing STP strength before and after training showed increases in subsets of neurons across low, intermediate, and high initialization conditions, with ten networks in each condition (Figure 2c; Appendix A.2): the networks learned to use stronger STP while learning the task. STP recruitment and higher activity-silent prevalence persisted across initializations and physiological parameter settings (Appendix A.6).

Because experiments report both single-trial stepping and stimulus-aligned ramping [Latimer et al., 2015], we asked whether transition-time variability across trials reproduces their coexistence in the trained networks. Transition times varied across trials, so stimulus-aligned averaging produced a gradual ramp whereas transition-aligned averaging recovered the abrupt rise (Figure 2d), and stronger evidence advanced the transition whereas weaker evidence prolonged the low-activity period (Figure 2e).

We then tested whether the distinction holds at the population level, using the choice output (the target readout minus the nontarget readout on correct-response trials). Step-like and ramp-like trajectories coexisted in both conditions, but step-like trajectories were more prevalent in RNNs with STP for every training seed (0.587 ± 0.040 versus 0.383± 0.036 across networks, mean ± SD; Figure 2f). Fitting procedures and numerical summaries are given in Appendices A.3 and A.5.

Setting 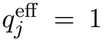 with trained weights held fixed reduced final-choice accuracy by 12.21 percentage points in trained RNNs with STP (Figure 2g). The decrease was larger on weak-evidence trials, supporting a contribution of STP-mediated transmission to decision performance (Appendix A.7).

## 4 Synaptic evidence and choice-selective circuit organization

Having established a behavioral contribution of STP, we next analyze synaptic evidence representations, neuronal activity, and connectivity in the trained networks to understand the mechanism underlying their activity-silent decision-making.

### 4.1 Synaptic evidence and the contribution of choice-selective populations

To examine how STP contributes to evidence accumulation, we compared linear readouts of cumulative sensory input from synaptic states and neuronal activity. Effective synaptic efficacy 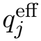 provided a more accurate linear readout of accumulated sensory evidence than neuronal activity *r* (Figure 3a). This advantage was consistent across all ten networks (Appendix A.8).

**Figure 3:**
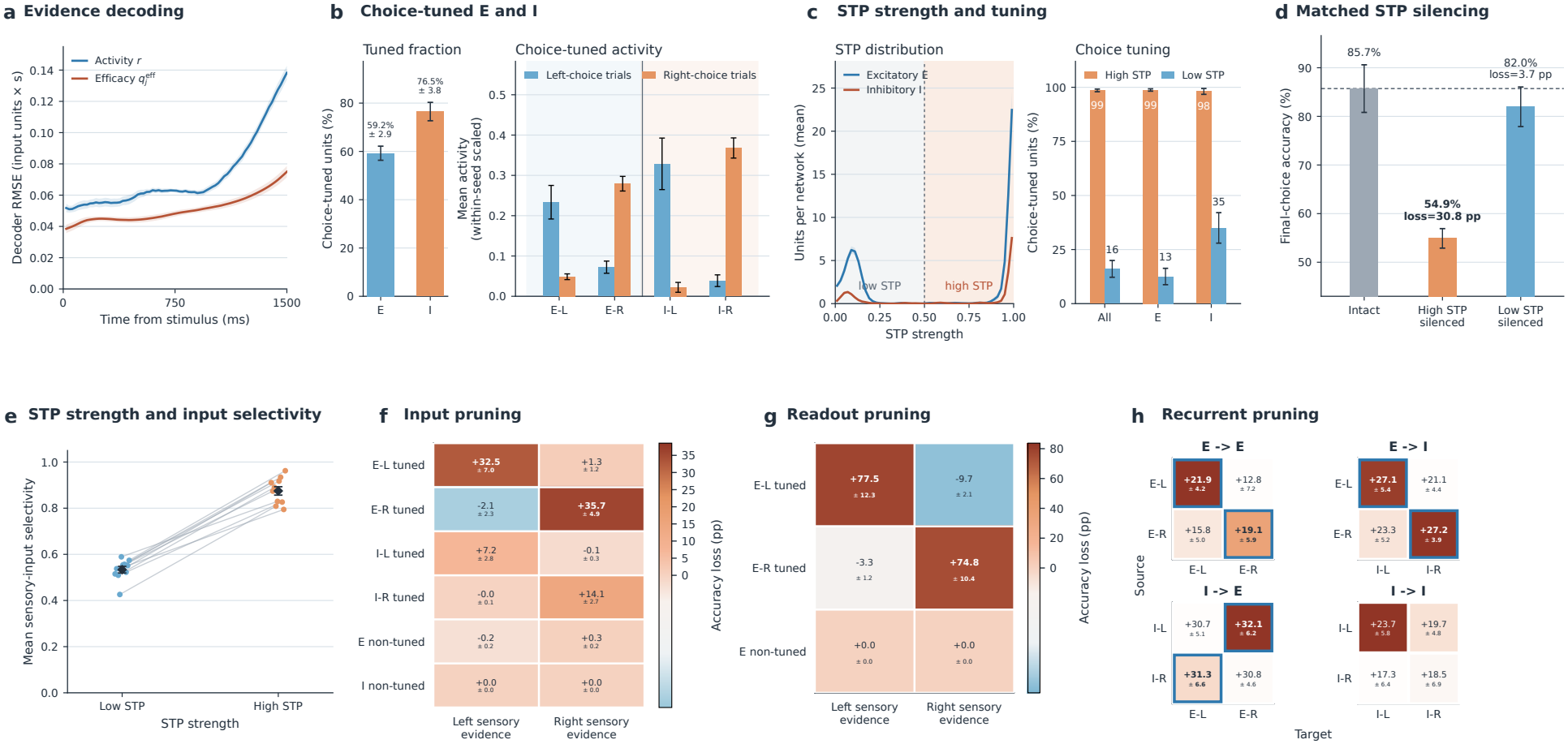
Synaptic evidence decoding, choice tuning, and circuit organization. **a**, Held-out RMSE for cumulative sensory-evidence decoding from *r* and 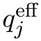 across time; lower values indicate more accurate decoding. A single decoder per representation is used across time. Curves and shading show mean ± SEM across ten networks. **b**, Fractions of choice-selective excitatory and inhibitory neurons and their population activity. **c**, Distributions of learned STP strength and choice-selective fractions among neurons with high or low STP strength. **d**, Neuronal contributions to decision performance, assessed by silencing high- or low-STP neurons with matched neuron counts and E/I composition. **e**, Mean normalized sensory-input selectivity in low- and high-STP groups. Connected points show paired network means; diamonds and error bars show mean ± SEM across ten networks. **f**, Accuracy loss after removing a sensory-input channel’s connections to each population. **g**, Accuracy loss for each evidence direction after disconnecting an excitatory population from both outputs. **h**, Accuracy loss after removing all recurrent connections from one choice-selective population to another; outlines mark selected pathways with large mean effects. Error bars and matrix labels report the mean ± standard error of the mean (SEM) across ten networks. Analysis details are given in Appendices A.8–A.12.

We next identified choice-selective neurons whose activity differed significantly between correct left and right choices (Appendix A.9). We found that both the excitatory and inhibitory populations contained substantial fractions of neurons preferring each choice (59.2 ± 2.9% of excitatory and 76.5 ± 3.8% of inhibitory neurons, mean ± SEM; Figure 3b). We denote the left- and right-preferring excitatory populations by *E*_*L*_ and *E*_*R*_, and the inhibitory populations by *I*_*L*_ and *I*_*R*_.

We then asked whether this choice selectivity was associated with the learned STP strengths. As Figure 2c showed, training increased STP strength in only a subset of neurons, splitting the learned strengths into a high group converging to values near one and a low group maintaining weak STP (Figure 3c). At the minimum-validation-loss checkpoints, high-STP neurons constituted 56.1 ± 10.5% of each network (mean ± SD). Separating neurons into high-STP and low-STP groups (Appendix A.10), we found that almost all high-STP neurons were choice-selective, whereas choice selectivity was much less common among low-STP neurons (Figure 3c).

To test the functional role of the predominantly choice-selective high-STP population, we silenced high-STP or low-STP neurons while matching their number and E/I composition. Silencing high-STP neurons reduced accuracy to near chance, whereas silencing low-STP neurons had only a small effect (85.7% intact, 54.9% after high-STP silencing, and 82.0% after low-STP silencing; Figure 3d). These results identify the high-STP, choice-selective population as an important contributor to evidence-guided decisions.

### 4.2 Connectivity of choice-selective populations

The association between STP strength and choice selectivity led us to examine the network’s connections. Neurons with stronger STP received more selective left–right sensory input, with a positive association in all ten networks (Figure 3e; Appendix A.11).

We further explored the circuit connectivity organization of the trained RNNs. The connection analyses use *connection pruning*, which removes selected connections by setting their weights to zero and measures the resulting change in task accuracy.

We first tested whether each choice-selective population depended on sensory evidence for its preferred choice. We found that left-preferring populations depended more strongly on the left-evidence input, while right-preferring populations depended more strongly on the right-evidence input (Figure 3f). Removing sensory connections to nonselective populations had much smaller effects. We then tested how the populations influenced the network’s output. Disconnecting *E*_*L*_ from both outputs reduced accuracy on left-evidence trials and increased accuracy on right-evidence trials; disconnecting *E*_*R*_ produced the reverse pattern (Figure 3g). Disconnecting nonselective excitatory neurons had little effect. These results show that each choice-selective population preferentially receives evidence for its preferred choice and that each choice-selective excitatory population contributes to the output favoring that choice.

We then examined how recurrent connections allow these populations to interact (Appendix A.12). Our connection-pruning analysis identified a pattern of self-excitation and mutual inhibition (Figure 3h). Removing excitation within a choice-selective population impaired accuracy more than removing excitation between the two populations. Removing an excitatory population’s projections to inhibitory neurons with the same choice preference caused a larger accuracy loss than removing its projections to inhibitory neurons with the opposite preference. Removing inhibitory projections to the opposite-choice excitatory population caused large accuracy losses as well. Together, these effects support an organization in which each excitatory population reinforces its own activity and recruits inhibitory neurons that suppress the competing excitatory population.

The excitatory-to-inhibitory and inhibitory-to-excitatory pathways resemble connections observed in mouse posterior parietal cortex: excitatory neurons preferentially connect to inhibitory neurons with the same choice preference, and inhibitory neurons preferentially connect to excitatory neurons with the opposite preference [Kuan et al., 2024].

## 5 theoretical model of activity-silent decision dynamics

The trained RNNs motivate our theoretical model through two observations: STP promotes activity-silent decision dynamics and supports evidence accumulation, and pruning reveals a choice-selective E/I circuit with self-excitation and mutual inhibition. Guided by these observations, we construct an independent four-population model with two competing choice groups, each comprising an excitatory and an inhibitory population (*E*_*L*_, *I*_*L*_, *E*_*R*_, and *I*_*R*_; Figure 4a). The model combines the self-excitation and mutual inhibition of classical decision circuits [Wang, 2002, Wong and Wang, 2006] with population-level STP. The model tests how evolving synaptic efficacy can support evidence accumulation and trigger a rapid choice transition by reshaping neuronal stability.

**Figure 4:**
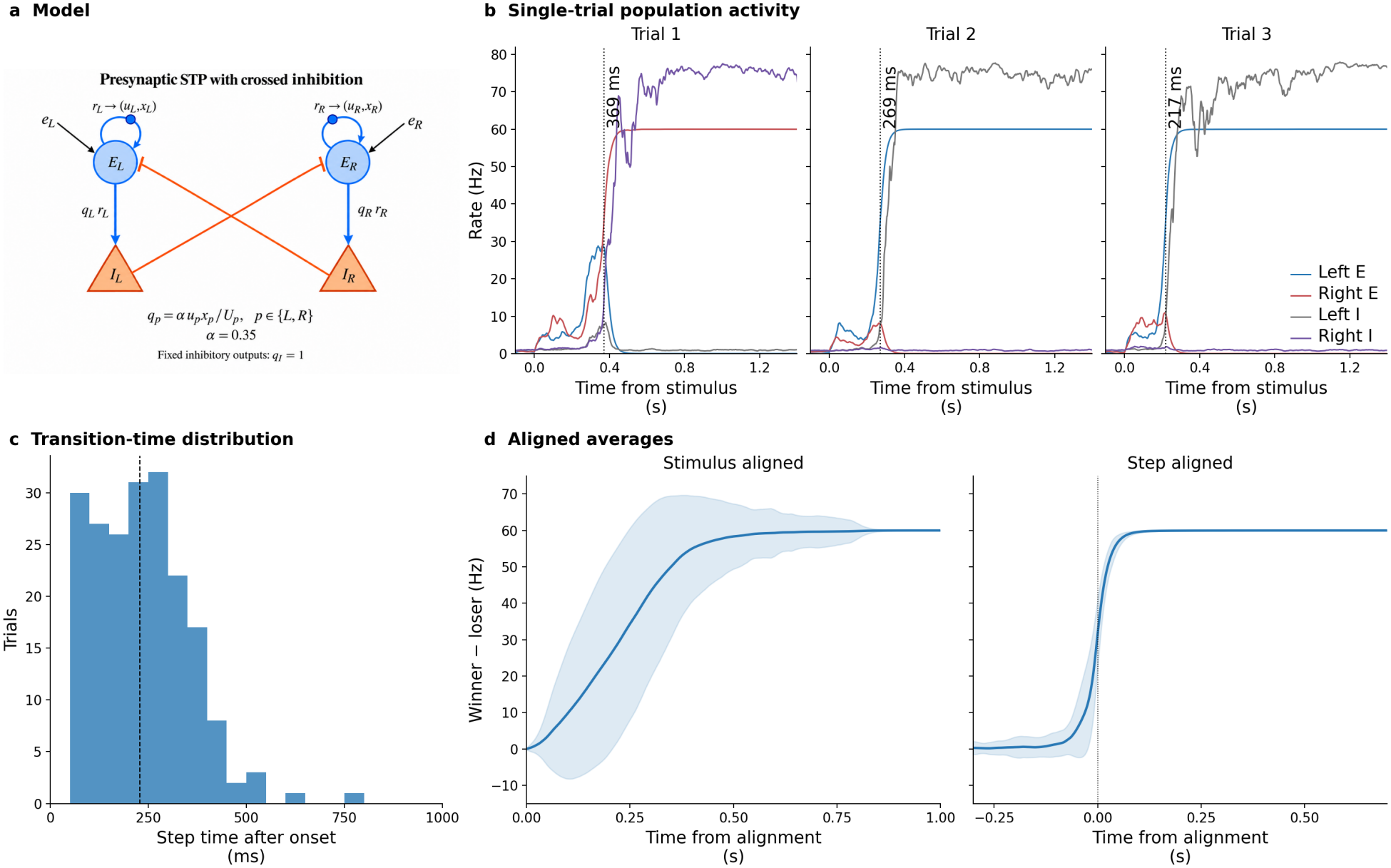
Activity-silent decision dynamics in the four-population model. **a**, Each excitatory population excites itself and its same-choice inhibitory population, which inhibits the opposite excitatory population. STP acts on excitatory outputs; inhibitory outputs are static. **b**, The first three example trials show the activity of all four populations. Dashed lines mark fitted transition times. **c**, Distribution of detected transition times; all 200 trials met the detection criteria. **d**, The same 200 trials averaged after alignment to stimulus onset or transition time; curves and shading show the mean and trial standard deviation (SD) of winner-minus-loser excitatory activity. Transition detection is described in Appendix B.2.

### 5.1 Model

For each choice *p* ∈ {*L, R* , *r*_*E,p*_ *}* and *r*_*I,p*_ denote the mean firing rates (in Hz) of the excitatory and inhibitory populations representing that choice, respectively. The index 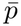 denotes the opposite choice. These firing rates evolve according to

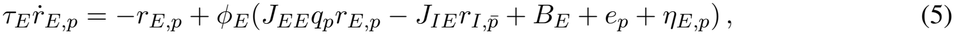

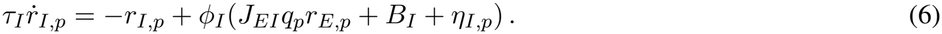

Here, *J*_*EE*_, *J*_*EI*_, and *J*_*IE*_ are positive connection strengths; *B*_*E*_ and *B*_*I*_ are background inputs; *e*_*p*_ is sensory evidence; and *η*_*E,p*_ and *η*_*I,p*_ are input noise. The activation functions *ϕ*_*E*_ and *ϕ*_*I*_ convert input into firing rate. To simplify simulation and analysis, we apply STP only to excitatory outputs, although adding STP to inhibitory outputs does not qualitatively change the results (Appendix B.1). The factor *q*_*p*_ = *αu*_*p*_*x*_*p*_*/U* represents the combined effect of STP on excitatory synaptic transmission, where *u*_*p*_ is synaptic utilization and *x*_*p*_ is the fraction of available synaptic resources. Each excitatory population has its own *u*_*p*_ and *x*_*p*_, and the resulting *q*_*p*_ scales both its self-excitation and its output to the same-choice inhibitory population. The synaptic variables evolve according to

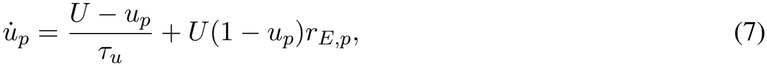

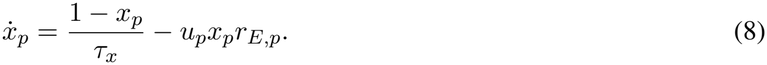

These equations describe activity-dependent facilitation and resource depletion [Masse et al., 2019]. The parameters *U* , *τ*_*u*_, and *τ*_*x*_ set baseline utilization and recovery times. The coefficient *α* = 0.35 scales excitatory synaptic transmission. Appendix B.1 specifies the activation functions, parameters, noise, and numerical implementation.

### 5.2 Activity-silent dynamics and their mechanism

Our simulations show that the model produces activity-silent decision dynamics on individual trials. After stimulus onset, both excitatory populations initially show low activity. One population then rises rapidly, recruits its same-choice inhibitory population, and suppresses the opposite excitatory population (Figure 4b). The transition time varies across trials (Figure 4c). Averaging after stimulus alignment therefore produces a gradual rise, whereas averaging the same trials after transition alignment preserves the rapid rise (Figure 4d).

To explain the delayed transition, we examine the Static network with *q*_*L*_ = *q*_*R*_ = *q* held fixed [Wong and Wang, 2006]. Increasing synaptic efficacy strengthens excitatory self-excitation and recruitment of the inhibitory populations. We calculate the equilibrium states and their stability to determine how these changes affect competition between the two choice groups (Figure 5a,b). The frozen-synapse analysis provides a quasi-static interpretation of how the current synaptic state reshapes neuronal stability, motivated by the slower dynamics of facilitation. The reported population dynamics are simulated in the full model with continuously evolving STP (Appendices B.2 and B.3).

**Figure 5:**
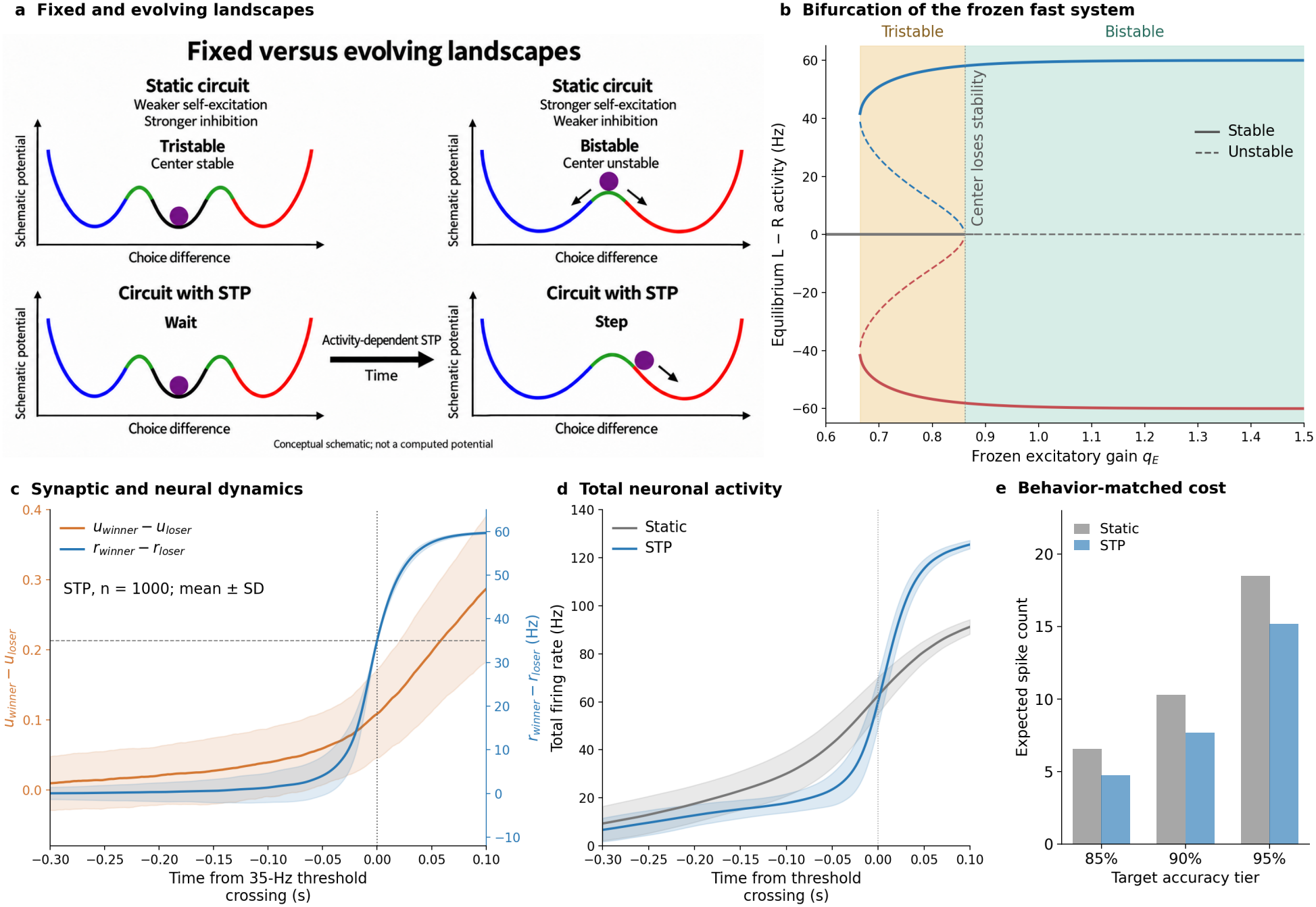
Synaptic mechanisms and activity cost. **a**, Conceptual phase portraits illustrating the transition from tristability to bistability as the central low-activity state loses stability; the curves are schematic, not computed from the model. **b**, Equilibrium excitatory activity differences with *q*_*L*_ = *q*_*R*_ = *q* held fixed over the displayed range *q* = 0.6–1.5. Solid and dashed lines denote stable and unstable equilibria, determined from the four-population Jacobian. The pale orange and pale green backgrounds mark the tristable and bistable regimes, respectively. The sampled boundaries are approximately *q* = 0.664 and *q* = 0.862; the unshaded interval to the left is monostable. **c**, Synaptic-utilization and excitatory-activity differences in the STP model, aligned to the first 35-Hz activity-difference threshold crossing (1000 trials; mean and trial SD). **d**, Sum of the firing rates of all four populations, aligned to the same decision threshold (1000 trials per condition; mean and trial SD). **e**, Behavior-matched activity-cost comparison at target-accuracy tiers of 85%, 90%, and 95%. Cost is the expected spike count integrated from stimulus onset to each model’s 35-Hz decision threshold. Bars show trial means at comparable accuracy and reaction time. Panel **d** retains the adjusted STP variant whose psychometric curve matches that of Static (Figure S6); Appendix B.4 gives its display protocol. The three-tier protocol for panel **e** is given in Appendix B.5.

Our stability analysis shows that, at intermediate synaptic efficacy, a stable low-activity state coexists with two stable choice states, forming a tristable regime. Further increases in synaptic efficacy destabilize the low-activity state, which becomes a saddle point, while the two stable choice states remain. This transition from tristability to bistability explains why a small activity difference can remain suppressed and then grow rapidly. The stability transition and activity-silent behavior persist over a finite grid of circuit and STP parameters (Appendix B.3).

Our simulations show that synaptic utilization begins to differ between the winning and losing populations before the rapid neuronal transition. When trials are aligned to the decision threshold, the difference in synaptic utilization between the winning and losing populations develops while their activity difference remains small (Figure 5c). These dynamics support the proposed role of synaptic states in retaining input history during the low-activity period: a linear decoder on the utilization and resource differences recovers evidence direction above chance throughout this period (Appendix B.2).

We next examine how the delayed rise in neuronal activity affects the expected spike count. Parameters of the STP model are adjusted so that its psychometric curve, which measures choice accuracy across evidence levels, matches that of the Static model; all other parameters, including the Static excitatory time constant, are unchanged. We measure total neuronal activity as the sum of the firing rates of all four populations. The STP model maintains lower total activity before the decision threshold than the Static model (Figure 5d). Integrating total firing rate from stimulus onset to threshold crossing gives the expected decision-period spike count. We compared this count across three target-accuracy tiers with comparable accuracy and reaction time (Figure 5e; Appendix B.5, Table 12). STP required fewer expected spikes at every tier, supporting an activity-cost advantage.

## 6 Discussion and conclusion

In summary, we trained excitatory–inhibitory RNNs with learnable STP on a reaction-time decision task and found that they developed activity-silent decision dynamics: choice-related activity remained low after stimulus onset and rose abruptly at transition times that varied across trials, so that stimulus-aligned trial averages reproduced ramping. Learning increased STP strength in a subset of neurons that were almost exclusively choice-selective and contributed most to accurate decisions. Fixed-weight STP interventions and decoding further link STP to decision performance and the representation of accumulated sensory evidence.

Connection pruning revealed a choice-selective E–I circuit of self-excitation and mutual inhibition. Guided by the choice-selective E/I organization revealed by pruning and STP’s contribution to activity-silent decisions, our theoretical model illustrates how evidence accumulation in synaptic efficacy can reshape neuronal stability and trigger a rapid choice transition.

Analysis of the trained RNNs identifies connections that can implement this competition, resembling the choice-dependent connections measured in posterior parietal cortex [Kuan et al., 2024] and relating to models in which inhibitory connectivity controls decision dynamics [Roach et al., 2023]; this mechanism complements experimental evidence for changes in neuronal dynamics during decision-making [Luo et al., 2025].

Why did task optimization produce activity-silent decision dynamics even though nothing in the objective prescribed them? The objective contained no term rewarding low activity or energy saving, yet gradient descent on the task objective discovered a mixed regime in which neurons with strong STP carry much of the decision computation. A network equipped with STP naturally favors storing short-timescale information in synaptic efficacy rather than in sustained neural activity: synaptic efficacy evolves on recovery time constants that are slow relative to neuronal dynamics and therefore well suited to retaining input history over hundreds of milliseconds, whereas rapidly fluctuating neural activity is better suited to fast operations such as the abrupt commitment to a choice. This division of labor is consistent with earlier STP network models of working memory, in which trained networks likewise stored information primarily in synaptic states [Mongillo et al., 2008, Masse et al., 2019, Kozachkov et al., 2022].

Delaying strong neuronal activity may additionally reduce the metabolic cost of making a choice: spikes and postsynaptic currents require energy to restore ionic gradients [Attwell and Laughlin, 2001, Howarth et al., 2012], and this cost constrains sustained cortical activity [Lennie, 2003]. Integrated firing rate provides a biologically motivated proxy for spike-related energy use. At comparable behavior level, STP reduces the expected spike count needed to reach a choice.

These observations also speak to artificial intelligence. Test-time-training architectures incorporate STP-like mechanisms by updating part of the network during inference [Ba et al., 2016, Sun et al., 2020], yet the computational advantages of storing moment-to-moment information in such fast synaptic states, rather than in recurrent neuron states, have received little systematic discussion. Our results provide a biological case study: task-trained networks recruit STP for evidence-guided decisions, and synaptic states provide a more accessible linear readout of accumulated sensory evidence than neuronal activity, suggesting STP-style mechanisms as a design principle for short-term memory in adaptive networks.

## Reproducibility statement

Appendix A specifies the task, RNN dynamics, numerical implementation, optimization, checkpoint selection, and analysis procedures used for the reported network results. Appendix B gives the parameters, simulation procedure, transition detection, stability calculations, and expected-spike-count analysis for the theoretical model. The corresponding sections identify the trial sets, training seeds, checkpoints, and aggregation units used for each figure where applicable.

## AI use statement

Large language models were used for research execution, including inspecting implementations, developing model and analysis code, configuring and monitoring experiments, interpreting recorded outputs, and preparing scientific figures. It was also used to edit and polish the writing and mathematical presentation. The methods and experiments resulting from these uses are described in the paper. The authors reviewed AI-assisted code and outputs, checked experimental claims against the underlying records. The authors take responsibility for the design, execution, interpretation, and final content of the work.

## Appendix

### A Implementation and functional analysis of the RNN model

The methods follow the order of the main text: task and network specification, optimization and check-point selection, paired-configuration summary, temporal-dynamics analysis, STP robustness, fixed-weight interventions, evidence decoding, functional population assignment, STP-strength analysis and neuronal silencing, connectivity correlations, and connection pruning.

#### A.1 Trial generation and behavioral readout

The reaction-time task has ten nonzero conditions formed by coherence *c* ∈ {1, 2, 4, 8, 16 } % and two latent directions. Gradient minibatches sample coherence and direction uniformly on non-catch trials. The official task sets the gradient-trial catch probability to 5*/*11; validation cycles over the ten nonzero conditions and one catch condition. Each 1100-trial validation set therefore contains 100 trials from every condition. Zero-coherence trials are used only for behavioral evaluation. Gradient, validation, and parameter-initialization random streams are separate.

The clean task inputs are zero during fixation. During the stimulus, the target evidence channel receives (1 + 3.2*c/*100)*/*2, the other evidence channel receives (1 *−* 3.2*c/*100)*/*2, and the third channel receives 1. The simulator adds a baseline of 0.2 and Gaussian input noise with time-step-dependent scaling to all three channels and rectifies the resulting inputs. The latent direction, rather than the noisy realized inputs, defines the target choice. On catch trials all three clean task inputs remain zero for 2000 ms.

The two linear readouts are trained against piecewise-constant targets. On a non-catch training or validation trial, both targets equal 0.2 during the 100-ms fixation period. The output mask is zero during the first 300 ms after stimulus onset, so this interval contributes no task loss. During the remaining 500-ms response interval, the target is 1.2 for the output corresponding to the latent direction and 0.2 for the other output. On a catch trial, both targets remain at 0.2 and both outputs are supervised throughout the 2000-ms trial. Padding beyond the end of a trial has zero mask weight.

Training and validation non-catch trials therefore last 900 ms: 100 ms of fixation followed by 800 ms of motion evidence. The held-out simulations used for Figures 2 and 3 follow the official test-time protocol instead, with 300 ms of fixation and 1500 ms of continuously presented motion evidence. This duration change affects only evaluation; it does not alter the learned parameters or the 300-ms onset of response supervision used during training. Table 1 summarizes the training, validation, and evaluation protocols.

**Table 1:** Trial protocols used for training and the reported analyses. All held-out simulations use a 0.5-ms integration step and save outputs every 10 ms.

| Use | Fixation | Stimulus | Trials per network | Role |
|---|---|---|---|---|
| Gradient training | 100 ms | 800 ms | minibatches of 20 | Parameter optimization; catch probability 5/11 |
| Validation | 100 ms | 800 ms | 1100 per check | Checkpoint selection from ten evidence conditions and one catch condition |
| Figure 2 | 300 ms | 1500 ms | 2000 | Temporal-shape and transition-time analyses |
| Figure 3b,c | 300 ms | 1500 ms | 600 | Choice tuning at 16% coherence; panel c combines these labels with checkpoint STP strengths |
| Figure 3d,f–h | 300 ms | 1500 ms | 16,500 per condition | Accuracy under pruning or silencing |

**Table 2:**
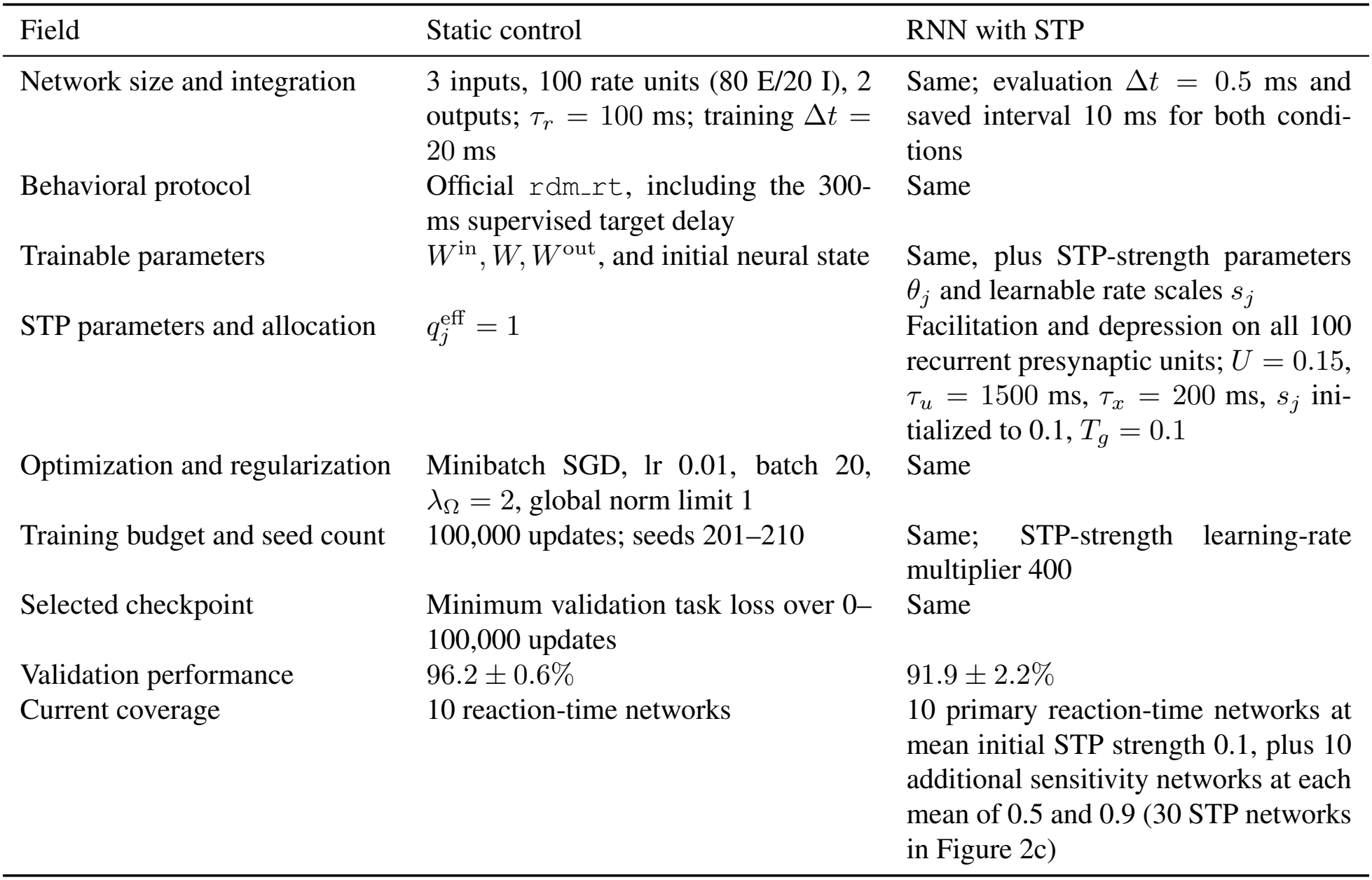
Implemented configuration and training coverage for the paired reaction-time comparison.

| Field | Static control | RNN with STP |
|---|---|---|
| Network size and integration | 3 inputs, 100 rate units (80 E/20 I), 2 outputs; τ<sub>r</sub> = 100 ms; training Δt = 20 ms | Same; evaluation Δt = 0.5 ms and saved interval 10 ms for both condi-tions |
| Behavioral protocol | Official rdm_rt, including the 300-ms supervised target delay | Same |
| Trainable parameters | W<sup>in</sup>, W, W<sup>out</sup>, and initial neural state | Same, plus STP-strength parameters θ<sub>j</sub> and learnable rate scales s<sub>j</sub> |
| STP parameters and allocation | q<sub>j</sub><sup>eff</sup> = 1 | Facilitation and depression on all 100 recurrent presynaptic units; U = 0.15, τ<sub>u</sub> = 1500 ms, τ<sub>x</sub> = 200 ms, s<sub>j</sub> ini-tialized to 0.1, T<sub>g</sub> = 0.1 |
| Optimization and regularization | Minibatch SGD, lr 0.01, batch 20, λ<sub>Ω</sub> = 2, global norm limit 1 | Same |
| Training budget and seed count | 100,000 updates; seeds 201–210 | Same; STP-strength learning-rate multiplier 400 |
| Selected checkpoint | Minimum validation task loss over 0–100,000 updates | Same |
| Validation performance | 96.2 ± 0.6% | 91.9 ± 2.2% |
| Current coverage | 10 reaction-time networks | 10 primary reaction-time networks at mean initial STP strength 0.1, plus 10 additional sensitivity networks at each mean of 0.5 and 0.9 (30 STP networks in Figure 2c) |

The two linear readouts are 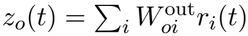 only excitatory neurons have nonzero output connections. Let *z*_*tbo*_ and *y*_*tbo*_ denote the predicted and target values for time point *t*, trial *b*, and output *o* ∈ *{*1, 2*}*, and let *m*_*tbo*_ ∈ *{*0, 1*}* denote the output mask. The task loss is the masked mean squared error

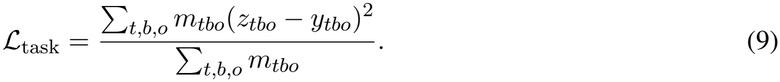

The normalization is over supervised output elements rather than first averaging within each trial or condition. The reported root-mean-squared error is 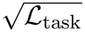.

Training and validation performance is the percentage of non-catch trials for which the larger output at trial end identifies the target choice. Threshold-based reaction time is an evaluation measure rather than the training objective. Outputs are sampled every 10 ms, and the response is the first saved sample at which the larger of the two linear outputs is strictly greater than 1.0; its argmax gives the reported choice. Thus the first crossing wins across samples, and the larger output resolves a same-sample double crossing. A trial is classified as a timeout if neither output exceeds 1.0 at any saved sample before the trial ends. For the held-out RNN trials, this deadline is 1800 ms from trial onset, or 1500 ms after stimulus onset. Reaction time is the crossing time minus stimulus-onset time. Analyses specify whether accuracy includes timeouts; latency summaries are restricted to the stated responded or correct-responded trials.

On the matched held-out streams, both conditions showed the expected dependence on sensory evidence: final-choice accuracy increased and median reaction time decreased with motion coherence (Figure S3). Accuracy is the final-output argmax over all attempted trials; reaction-time summaries include correct threshold responses only. Curves and error bars first summarize trials within each network and then report the mean and SEM across the ten independently trained networks.

#### A.2 Network dynamics and numerical implementation

Equations 1–4 define the neural and synaptic dynamics. We use *τ*_*r*_ = 100 ms, *U* = 0.15, *τ*_*u*_ = 1.5 s, *τ*_*x*_ = 0.2 s, an initial rate scale *s*_*j*_ = 0.1 s^*−*1^*/*a.u., *u*_*j*_(0) = *U* , *x*_*j*_(0) = 1, and *T*_*g*_ = 0.1 [Masse et al., 2019]. The initial conditions give *u*_*j*_(0)*x*_*j*_(0)*/U* = 1 and therefore preserve resting recurrent strength. The parameters and states (*a*_*j*_, *s*_*j*_, *u*_*j*_, *x*_*j*_) are shared by all recurrent projections leaving presynaptic neuron *j*; STP is not applied to the input or readout weights. The rate scale *s*_*j*_ is learned jointly with STP strength and the long-term weights. The learned parameters are held fixed during inference while *u*_*j*_ and *x*_*j*_ evolve within each trial.

The implementation combines two public code bases at fixed revisions:

- the task, E–I rate network, objective, optimizer, and long-term parameter initialization follow Pycog [Song et al., 2016] at revision `7c5d54b` (https://github.com/frsong/pycog);
- the utilization and resource-depletion updates are adapted from the code accompanying Masse et al. [2019], revision `6c3c486` (https://github.com/nmasse/Short-term-plasticity-RNN); and
- the normalized efficacy, learnable STP strength, truncated-Gaussian initialization of STP strength, and checkpoint snapshots are additions made within the Pycog training framework. No task, neuronal, readout, or long-term-weight component is imported from the STP source network.

The initial STP strengths are sampled from a normal distribution with mean *µ* and truncated to the open unit interval. We use *µ* ∈ {0.1, 0.5, 0.9} for the initialization comparison and *µ* = 0.1 for the primary multiseed analyses:

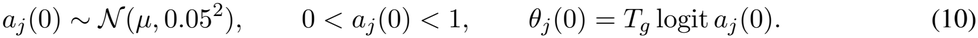

The STP strengths and rate scales *s*_*j*_ are learned jointly with the long-term weights; *U* , *τ*_*u*_, *τ*_*x*_, and the initial synaptic states remain fixed. The model uses no STP-strength regularizer. Unit-gain and depletion-off checks recover the corresponding static and facilitation-only computations to numerical precision.

For comparison with a Static network, resting effective recurrent strength is distinguished from the unmodulated weight matrix. When synaptic efficacy is normalized by its resting value, weight initialization and regularization use the same convention. The E/I constraint fixes the signs of outgoing recurrent weights.

All 100 recurrent units receive the three task inputs. Recurrent connectivity contains every directed connection except self-connections; Dale’s law fixes the sign of each unit’s outgoing recurrent weights. Input weights are nonnegative, and only the 80 excitatory units project to the two outputs. Recurrent weights are initialized from the official gamma distribution and rescaled to spectral radius 1.5; input and output weights use the official uniform initialization. Long-term weights (input, recurrent, and output) and the initial neural state are trainable. The initial state begins at *h*_*i*_(0) = 0.1 for every unit, whereas recurrent and output biases remain fixed at zero.

Input noise has simulator parameter *σ*_in_ = 0.01 and time constant 100 ms; recurrent noise has parameter *σ*_rec_ = 0.15. The simulator rescales both stochastic inputs with the integration step. Sensory inputs are rectified after adding the constant input level and input noise. Neural states are reset to the learned initial condition at each trial. Synaptic states are reset to *u*_*j*_(0) = *U* and *x*_*j*_(0) = 1 for every presynaptic neuron *j*.

The code evaluates the continuous dynamics using the forward Euler method. In the discrete equations below, Δ*t* and all time constants are expressed in milliseconds, and Δ*t*_*s*_ = Δ*t/*1000 is the time step in seconds. The superscript *n* denotes the integration step, and Π_[0,1]_ clips its argument to [0, 1]. One step is

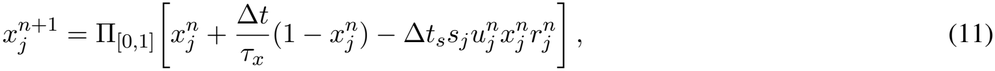

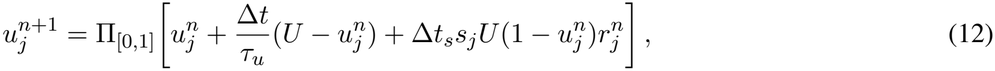

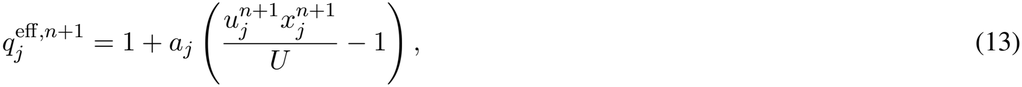

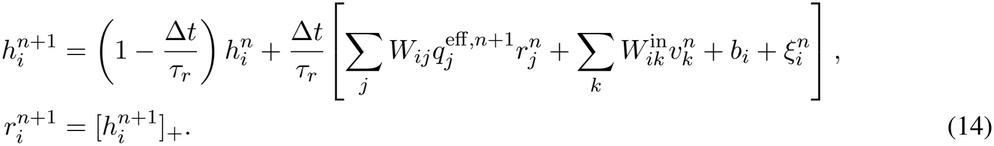

Thus the previous presynaptic rate first updates both synaptic states; their updated normalized product then modulates the recurrent drive, and the neural state is advanced last. Training uses Δ*t* = 20 ms. Held-out evaluation uses Δ*t* = 0.5 ms and saves activity and outputs every 10 ms while preserving the simulator’s time-step-dependent noise scaling.

#### A.3 Optimization and checkpoint selection

Minibatches contain 20 trials, and Equation 9 is evaluated jointly over all supervised elements in each minibatch. Optimization uses float32 stochastic gradient descent (SGD) with a base learning rate of 0.01 and global gradient-norm clipping at 1. The update includes the original vanishing-gradient correction *λ*_Ω_ ∇Ω, with *λ*_Ω_ = 2. All explicit L1 weight, L2 weight, activity-rate, and STP-strength penalties are zero. The logged task loss and regularized objective are therefore identical; the separately logged Ω value contributes to the parameter update through its gradient but is not added to the displayed objective.

Both conditions train for exactly 100,000 updates and validate every 500 updates on 1100 trials. Training is divided into 0–50k and 50–100k stages; the continuation preserves the first-stage best checkpoint, and final selection compares it with every continuation checkpoint. Checkpoints are selected by the minimum pure validation task loss over the complete 0–100k record rather than by root-mean-squared error, validation performance, or threshold-based reaction time. Specifically, if *K*_*s*_ contains the preserved first-stage best checkpoint and every available continuation checkpoint for seed *s*, the analyzed update is

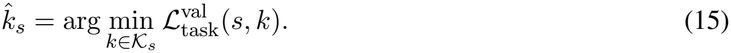

Static networks and RNNs with STP are selected by the same rule and from the same training budget.

For the RNNs with STP in Figures 2 and 3, only the raw STP-strength-parameter update is multiplied by 400; all other parameters retain the base learning rate. In the complete seed-201 training log, sampled every 500 updates from 500 to 100,000, the median raw *θ*-gradient norm was 7.39 *×*10^*−*5^ and the median full-parameter gradient norm was 9.64 *×*10^*−*2^. Their median ratio was 6.86 *×*10^*−*4^ (interquartile range, 4.08 *×*10^*−*4^–1.28 *×*10^*−*3^), placing the unscaled STP-strength-parameter gradient norm approximately three orders of magnitude below the full gradient norm.

A same-initialization, 100,000-update calibration in seed 201 compared multipliers of 200, 400, 500, 600, and 1000. Their minimum validation task losses were 0.06797, 0.06489, 0.06962, 0.07486, and 0.07722, respectively. All five runs remained finite, and 400 gave the lowest validation loss in this calibration, so this multiplier was fixed before the ten-network analysis.

RNNs with STP reached a validation performance of 91.9*±*2.2% at the selected checkpoints (mean*±*SD across ten networks), and their learning trajectories are shown in Figure 2a. Static networks reached 96.2 ± 0.6%, with learning trajectories shown in Figure S1. Validation performance uses the final-output argmax on the balanced non-catch validation trials and is distinct from threshold-response measures and the held-out test protocol. In the held-out comparison of ten STP and ten independently trained Static networks on the same 2,000-trial protocol, final-choice accuracy was 95.05 ± 1.17% and 79.92± 6.78%, respectively (mean SD across networks). Timeout fractions, defined in Appendix A.1, were 5.46 ±2.26% and 13.96 ± 3.26%, and correct threshold crossings within the trial occurred on 90.25 ±1.75% and 83.44 ±3.14% of trials.

**Figure S1:**
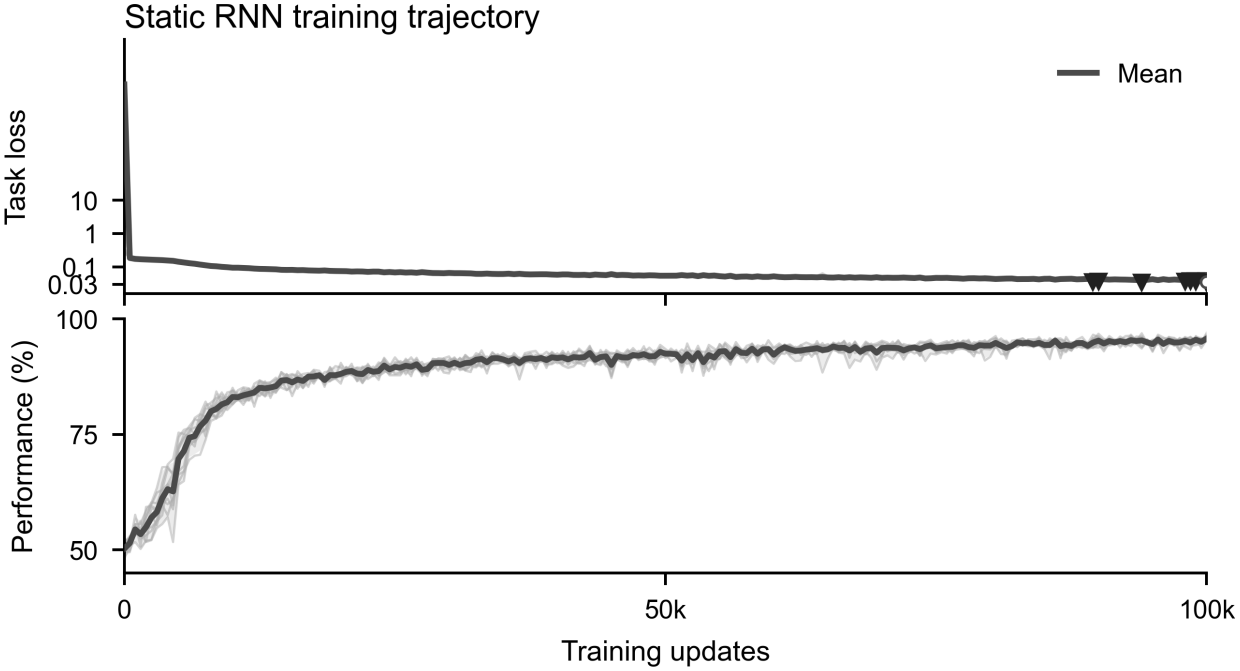
Static RNNs learn the reaction-time motion-discrimination task. Validation task loss and validation performance are shown over 100,000 training updates for ten independently initialized networks. Thin lines denote individual networks, the thick line denotes their mean, and shading shows the point-wise 2.5th–97.5th percentiles across networks. Downward triangles mark the minimum-validation-task-loss checkpoints; open circles mark the 100,000-update endpoints.

For training seed *s*, ordinary network parameters use seed *s*, while gradient and validation trials use fixed seeds 11 and 22, respectively. Static and STP runs with the same training seed share the ordinary initialization and task-stream seeds. For RNNs with STP, the Gaussian STP-strength sample uses seed *s* + 910000 and inverse-CDF truncation on (0, 1), avoiding point masses at the boundaries. Seeds 201– 210 define the ten primary STP networks paired with the ten Static networks. The initialization-sensitivity analysis contains ten additional networks initialized at mean STP strength 0.5 and ten initialized at 0.9, while retaining standard deviation 0.05, seeds 201–210, and the remaining protocol. Figure 2c therefore summarizes 30 STP trainings, ten per initialization condition; the 20 additional networks do not enter the primary paired comparisons.

For paired network-level comparisons, we use a two-sided exact paired randomization test that exhaustively flips the signs of the ten within-seed differences. Confidence intervals are percentile intervals obtained by resampling the ten paired networks 200,000 times with random seed 20260925. Trials and neurons within a network are not treated as independent replicates.

#### A.4 Configuration summary

The current main configuration and run inventory are summarized below. The RNN with STP is the default treatment, and the Static network is its computational control. Both retain the same reaction-time task, neural dynamics, long-term parameterization, optimizer, and training budget.

#### A.5 Ramp–step BIC comparison

##### A.5.1 Evaluation set and fitted models

Figure 2 is evaluated on 200 trials in each of the ten nonzero coherence–direction conditions (2000 trials per network), using random seed 100 and the 300-ms-fixation/1500-ms-stimulus test protocol. A trial batch is generated once and replayed through the paired Static RNN and RNN with STP, so their input and recurrent-noise realizations are matched; resetting the evaluation seed also gives every training seed the same external trial stream. Behavioral summaries include all attempted trials, whereas the shape analysis admits only trials with a correct threshold response and a complete requested window. The denominator of each displayed trajectory fraction is the number of eligible fitted trials in that network.

The network-level comparison in Figure 2f uses the target readout minus the nontarget readout in the response-aligned interval from *−*400 to 0 ms. Representative-neuron analyses in Figure 2b,d,e use the interval from 0 to 600 ms after stimulus onset. Both are sampled every 10 ms and fitted without smoothing.

For samples 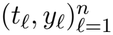 , the ramp model is 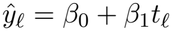 . The step model has one change point *k* and two constant levels,

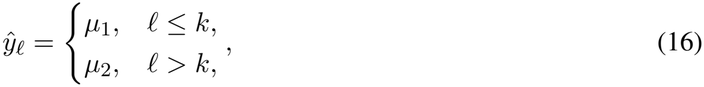

where every candidate leaves at least max(3, 0.1*n*) samples on each side; the candidate with minimum residual sum of squares is retained. Under a common independent-Gaussian residual approximation,

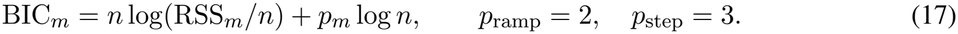

We define ΔBIC = BIC_ramp_ *−*BIC_step_ and label trials with ΔBIC *>* 2 as step-like, those with ΔBIC *< −*2 as ramp-like, and the remainder as indeterminate. This operational rule compares the fitted ramp and one-change-point step families.

Each trained network is the independent unit for condition-level comparisons. Trial fractions are first computed within each network, and paired differences then compare matched Static and STP seeds. Trials and units within a network are not treated as independent network replicates.

##### A.5.2 Neuron-level prevalence and pre-transition activity

We complemented the choice-output analysis with a split-sample neuronal analysis. Even-numbered trials determined each neuron’s preferred choice from its mean activity during the final 100 ms of the 0–600-ms fitting window. Odd-numbered correct threshold-response trials in that preferred direction were used for classification. We retained neurons with at least 20 assessment trials and peak activity of at least 0.2 on at least half of those trials. A retained neuron was classified as activity-silent when at least half of its active trials favored an upward step (ΔBIC *>* 2 and *µ*_2_ *> µ*_1_) and its median normalized pre-transition deviation across those upward-step trials was at most 0.2. For trial *i*, this deviation was

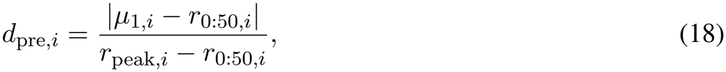

where *r*_0:50,*i*_ is mean activity during the first 50 ms after stimulus onset. This permits a positive resting rate but requires the fitted pre-transition level to remain close to the trial’s own initial activity.

Across the ten RNNs with STP, 32.9 ± 9.7% of recurrent neurons met this joint shape-and-pre-transition-activity criterion (mean ± SD), compared with 18.1 ±4.7% in the ten paired Static RNNs. The paired mean increase was 0.148 (95% bootstrap confidence interval, 0.089–0.210; exact paired-randomization *p* = 0.004). The corresponding excitatory and inhibitory fractions were 34.8 ± 9.1% and 25.5 ± 15.9% in RNNs with STP, and 16.8 ± 3.8% and 23.5 ±11.6% in Static RNNs. The choice-output trajectory fraction in Figure 2f instead quantifies trajectory shape alone using ΔBIC *>* 2; it does not apply the neuron-level pre-transition-activity criterion.

The main-text prevalence restricts the denominator to choice-tuned neurons. Let *A*_*s*_ be the activity-silent neurons identified above and *C*_*s*_ the choice-tuned neurons in network *s*, classified by the procedure in Appendix A.9. Choice tuning was evaluated independently at the selected checkpoint of each Static and STP network using the same high-coherence trials, analysis window, and Welch-test/BH-FDR rule. For each network, we computed

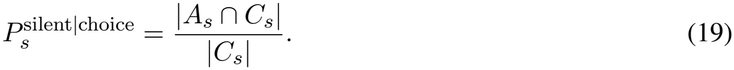

The ten network-level fractions were then averaged with equal weight. They were 52.2 ± 16.2% for RNNs with STP and 36.2 ±7.0% for Static RNNs (mean ±SD). The paired mean increase was 0.160 (95% bootstrap confidence interval, 0.068–0.249; exact paired-randomization *p* = 0.016). Conversely, 97.6 ±4.4% of activity-silent neurons in RNNs with STP and 98.2 ±3.5% in Static RNNs were choice tuned.

For the neurons meeting the joint criterion, mean activity during the final 50 ms before the fitted transition was 41.2 ±2.2% of the raw trial peak. After subtracting each trial’s initial activity and normalizing by its initial-to-peak range, the pre-transition deviation was 23.8 ±0.9%. These summaries are distinct from the fitted-level threshold in Equation 18 and provide a complementary description of the low-activity period.

##### A.5.3 Representative units and display conventions

Representative units for Figures 2d,e are selected on a separate discovery set containing 200 trials per direction at 16% coherence. Selection is restricted to excitatory units whose absolute readout-weight difference is at least the E-unit median, whose peak activity reaches 0.2 on at least half of valid preferred-direction trials, and whose stimulus-aligned trials are classified as step-like at least half of the time. The unit closest to the eligible subgroup median in step fraction and peak activity is retained. Static representative units use the analogous majority-ramp criterion. These units are then evaluated on the independent 2000-trial stream described above.

Figure S2 illustrates the scoring rule on the four paired seed-210 trials shown in Figure 2b. Static unit 68 is paired with STP units 52, 32, 57, and 32 on trial IDs 1319, 1209, 1589, and 1611. The STP units passed the independent discovery screen; one clear paired example was then chosen from each of four fixed transition-time windows (50–100, 120–180, 430–480, and 220–270 ms).

For Figure 2b, each trajectory is scaled by its 5th and 95th activity percentiles and clipped to [0, 1]. A centered 30-ms moving average is used only for the colored display line; the raw trace and all fits remain unsmoothed. Figure 2d uses 171 correct, preferred-direction trials from the representative STP unit at 16% coherence, restricted to step-classified trials with change points between 50 and 450 ms. The same percentile-normalized trials are averaged after alignment to stimulus onset or fitted change point. Bands are 95% percentile intervals from 5000 trial-level bootstrap resamples and quantify within-network uncertainty. Figure 2e groups within-network median transition times as low (1–2%), medium (4–8%), or high (16%) coherence and reports the mean and SD across networks. Figure 2c averages network-level empirical cumulative distribution functions (ECDFs) of the 100 STP strengths and shows pointwise 2.5th–97.5th percentiles across networks.

Across networks, the mean within-network median transition time decreased from 162 ±28 ms at low coherence to 115 ±10 ms at high coherence (mean ±SD). Applying the same change-point procedure to the independently selected majority-ramp Static representative units gave 272 ±98 ms at low coherence, 233 ±88 ms at medium coherence, and 183 ±61 ms at high coherence. Nine Static networks contributed a low-coherence median because one had no eligible low-coherence step-classified trial; all ten contributed to the other groups. Activity-silent, step-like choice-output trajectories, operationally defined by ΔBIC *>* 2, were more prevalent with STP. Their fractions were 0.587 ±0.040 for RNNs with STP and 0.383 ±0.036 for Static networks. The paired mean increase was 0.204 (95% bootstrap confidence interval, 0.177–0.231; exact paired-randomization *p* = 0.002). The corresponding ramp-like fractions were 0.350 *±* 0.037 and 0.579 *±* 0.033, and indeterminate fractions were 0.063 *±* 0.004 and 0.038 *±* 0.008.

**Figure S2:**
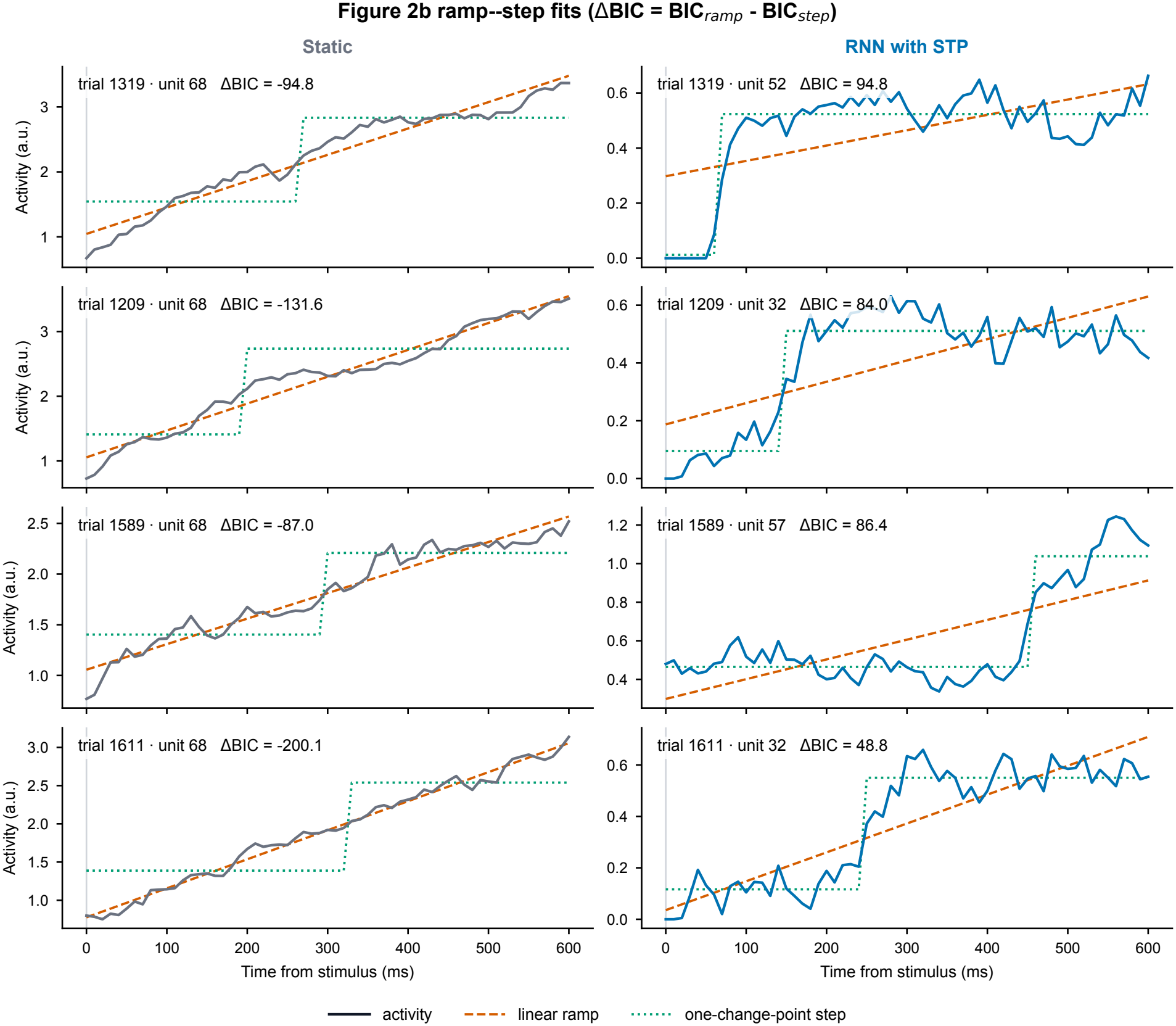
Representative fits for the ramp–step BIC comparison. Rows are the same four paired seed-210 trials shown in Figure 2b; columns show the Static and STP networks. Solid curves show raw unsmoothed activity from 0 to 600 ms after stimulus onset, dashed orange curves show fitted ramps, and dotted green curves show fitted one-change-point steps. Unit identifiers and ΔBIC values are given in each panel. Positive ΔBIC favors the step model and negative values favor the ramp model.

**Figure S3:**
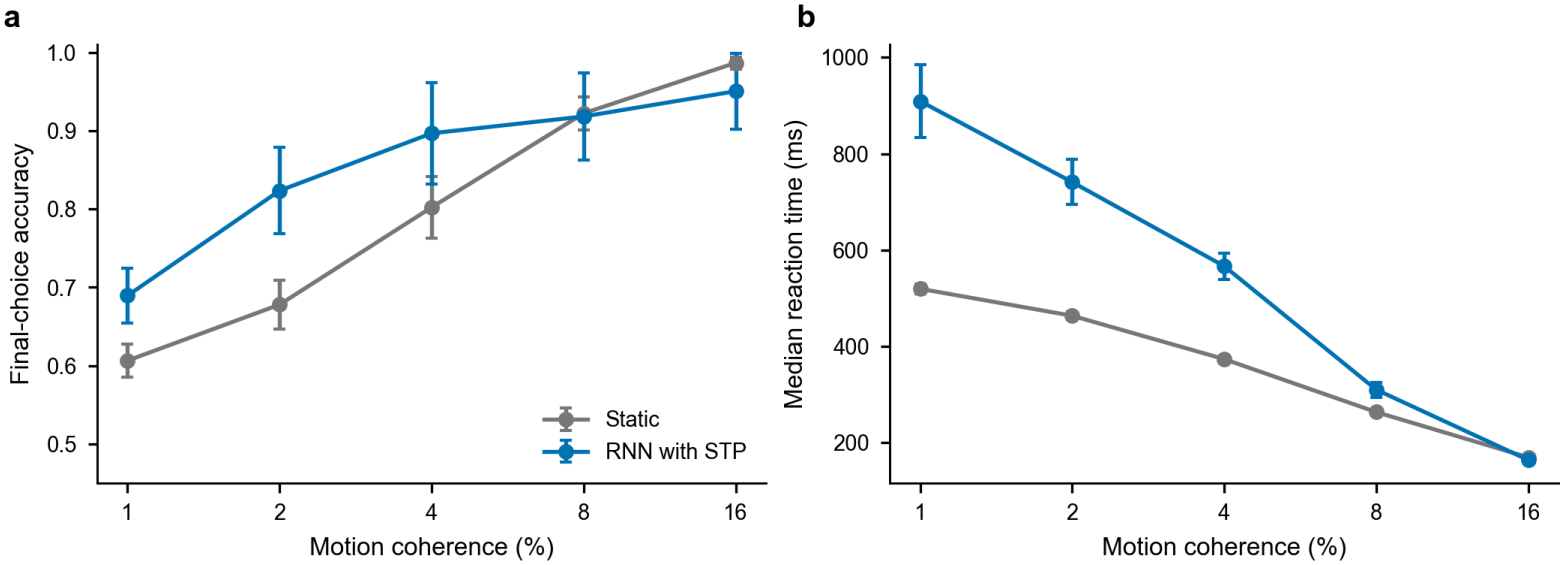
Held-out behavior across motion coherence. **a**, Final-choice accuracy over all attempted nonzero-coherence trials. **b**, Median reaction time on correct threshold-response trials. Points and error bars show the mean and SEM across ten paired networks; each network contributes 400 trials per coherence, equally divided between the two directions.

#### A.6 STP sensitivity and robustness tests

We retrained ten networks under each of nine conditions (90 networks total). Initial STP strengths were sampled with standard deviation 0.05 around means of 0.1, 0.5, or 0.9. The six baseline-parameter conditions varied utilization *U* , facilitation recovery time *τ*_*u*_, or resource recovery time *τ*_*x*_ one at a time by a factor of 0.5 or 2. Training lasted 100,000 updates, with minimum-validation-task-loss checkpoint selection and the remaining settings from Appendix A.3.

Output activity-silent fractions use target-minus-nontarget output in the 400-ms response-aligned window and the BIC rule in Appendix A.5. STP and same-seed Static networks share common-valid trials: correct threshold responses with a complete fitting window. Strength change is checkpoint minus initial network-mean strength; fraction change is the paired STP-minus-Static difference. We also computed within-network Spearman correlations between STP strength and sensory-input strength (*ρ*_input_), defined as the *L*_2_ norm of the two sensory-input weights, or the binary choice-tuned label (*ρ*_choice_). Choice tuning used the 1400–1800-ms activity window on correct threshold responses at 16% coherence, with 300 generated trials per direction and Welch tests corrected by BH-FDR at *q <* 0.05.

Across all nine conditions, STP strength increased in all 90 networks. For the three initialization means, observed network-mean strengths increased from 0.104 to 0.587, from 0.501 to 0.697, and from 0.898 to 0.938, respectively. Output activity-silent fractions exceeded the matched Static controls in all ten pairs in every condition (two-sided exact paired sign-flip *p* = 0.00195 per condition). Mean validation final-choice accuracy ranged from 89.04% to 95.14% across the six baseline-parameter conditions. Both *ρ*_input_ and *ρ*_choice_ were positive in all 90 networks; their condition means ranged from 0.602 to 0.734 and from 0.547 to 0.716, respectively. Tables 3 and 4 summarize all metrics, with strength changes, fractions, and accuracy reported as mean*±*SEM and correlations as mean*±*SD across ten networks per condition.

**Table 3:** STP-strength initialization sensitivity. Strength changes and fractions are mean*±*SEM; correlations are mean*±*SD (ten networks per condition). Fraction changes are relative to matched Static controls.

| Initial mean | Strength increase | STP fraction | Static fraction | Paired increase | ρ<sub>input</sub> | ρ<sub>choice</sub> |
|---|---|---|---|---|---|---|
| μ = 0.1 | 0.484 ± 0.030 | 0.586 ± 0.015 | 0.401 ± 0.016 | 0.184 ± 0.022 | 0.635 ± 0.060 | 0.699 ± 0.095 |
| μ = 0.5 | 0.196 ± 0.023 | 0.625 ± 0.012 | 0.409 ± 0.019 | 0.215 ± 0.024 | 0.633 ± 0.126 | 0.648 ± 0.146 |
| μ = 0.9 | 0.040 ± 0.004 | 0.568 ± 0.014 | 0.409 ± 0.016 | 0.159 ± 0.023 | 0.618 ± 0.054 | 0.716 ± 0.069 |

**Table 4:** Sensitivity to baseline STP parameters. Strength changes, paired fraction increases, and accuracy are mean *±* SEM; correlations are mean *±* SD (ten networks per condition). Fraction increases are relative to matched Static controls.

| Condition | Strength increase | Paired fraction increase | Validation accuracy | ρ<sub>input</sub> | ρ<sub>choice</sub> |
|---|---|---|---|---|---|
| *U* × 0.5 | 0.422 ± 0.031 | 0.314 ± 0.020 | 89.04 ± 0.32% | 0.734 ± 0.118 | 0.547 ± 0.148 |
| *U* × 2 | 0.416 ± 0.016 | 0.084 ± 0.017 | 95.14 ± 0.36% | 0.697 ± 0.064 | 0.693 ± 0.068 |
| τ<sub>*u*</sub> × 0.5 | 0.517 ± 0.016 | 0.210 ± 0.027 | 92.72 ± 0.62% | 0.602 ± 0.076 | 0.674 ± 0.113 |
| τ<sub>*u*</sub> × 2 | 0.485 ± 0.033 | 0.219 ± 0.016 | 91.93 ± 0.59% | 0.624 ± 0.062 | 0.704 ± 0.099 |
| τ<sub>*x*</sub> × 0.5 | 0.494 ± 0.026 | 0.181 ± 0.024 | 92.62 ± 0.65% | 0.639 ± 0.044 | 0.714 ± 0.070 |
| τ<sub>*x*</sub> × 2 | 0.480 ± 0.030 | 0.190 ± 0.023 | 92.86 ± 0.58% | 0.618 ± 0.074 | 0.664 ± 0.156 |

#### A.7 Fixed-weight interventions and component ablations

We analyzed ten trained STP networks (seeds 201–210) at their minimum-validation-task-loss checkpoints. These analyses examine how STP strength relates to trained connectivity, how STP interventions affect the fixed circuit, and how synaptic states encode accumulated input. The independently trained network is the unit of replication; trials and neurons are nested observations. Evaluation streams are specified separately for each analysis.

We replayed each selected network with its trained input, recurrent, and output weights, external inputs, and trial noise held fixed. Interventions set effective synaptic efficacy to one 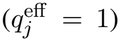 , retain facilitation alone (*x* = 1), or retain depression alone (*u* = *U*). Final-choice accuracy is measured on the same 2,000 nonzero-coherence trials per network (evaluation seed 100). All ten networks contribute to the accuracy summaries, including networks whose facilitation-only outputs become extreme. Table 5 reports mean*±*SEM and paired intervention-minus-intact changes.

**Table 5:** Fixed-weight STP interventions. Accuracy uses final-output argmax across all ten networks; changes are paired within network.

| Condition | Final-choice accuracy | Paired change (points) |
|---|---|---|
| Intact STP | 94.34 ± 0.74% | Reference |
| Effective synaptic efficacy set to one | 82.13 ± 3.35% | −12.21 ± 3.69 |
| Facilitation only (depression removed) | 57.79 ± 4.29% | −36.55 ± 4.14 |
| Depression only (facilitation removed) | 82.58 ± 2.46% | −11.76 ± 2.77 |

Setting efficacy to one reduced accuracy by 12.21 percentage points (95% CI for the paired change, *−*20.55 to *−*3.87; exact paired sign-flip *p* = 0.00586). On weak-evidence trials (absolute coherence 1 or 2), the mean decrease was 19.40 points. Neither component alone preserved intact performance. These interventions show a causal contribution of STP-mediated recurrent transmission to decision performance.

##### Amplitude-normalized neuronal transitions

We fitted *r*(*t*) = *b* + *A* sigmoid((*t − t*_0_)*/k*) to high-STP neurons (strength *≥*0.5) preferring the intact network’s actual threshold choice, using 10-ms samples during the first 600 ms after stimulus onset. The raw midpoint slope *A/*(4*k*) depends on both amplitude and direction. We therefore report amplitude-normalized absolute slope, 1*/*(4*k*), and the 10–90% transition width, 2 log(9)*k* (Table 6). For slopes, *k* is expressed in seconds; widths are reported in milliseconds.

**Table 6:**
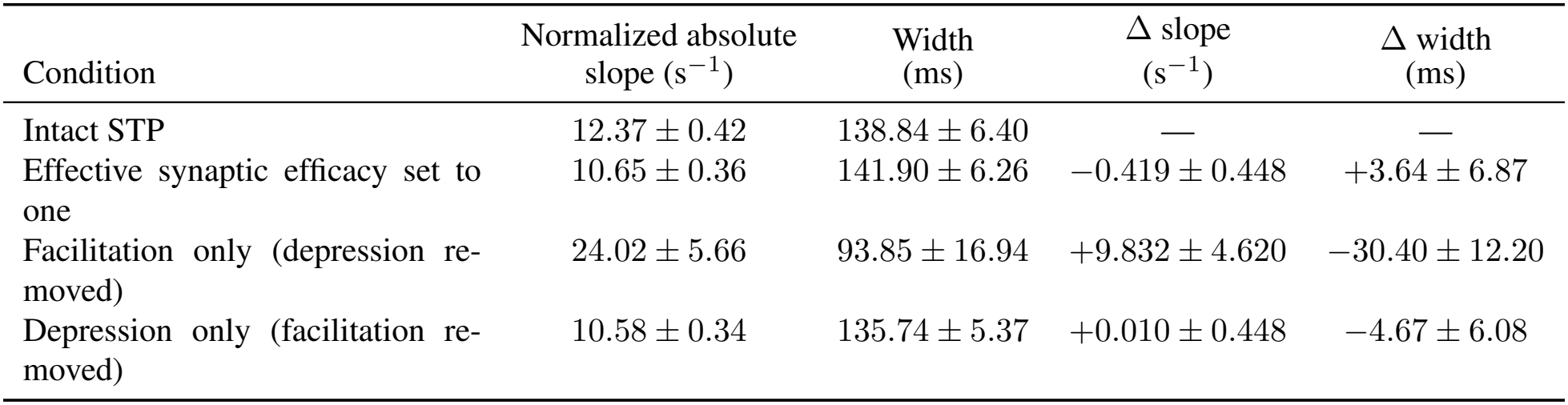
Amplitude-independent transition measures across all ten networks (mean ±SEM). Absolute values summarize each condition’s accepted fits; Δ values summarize within-network medians of paired intervention-minus-intact changes using the same trials and neurons. These paired changes need not equal differences between condition means.

| Condition | Normalized absolute<br>slope (s<sup>−1</sup>) | Width<br>(ms) | Δ slope<br>(s<sup>−1</sup>) | Δ width<br>(ms) |
|---|---|---|---|---|
| Intact STP | 12.37 ± 0.42 | 138.84 ± 6.40 | — | — |
| Effective synaptic efficacy set to one | 10.65 ± 0.36 | 141.90 ± 6.26 | −0.419 ± 0.448 | +3.64 ± 6.87 |
| Facilitation only (depression re-moved) | 24.02 ± 5.66 | 93.85 ± 16.94 | +9.832 ± 4.620 | −30.40 ± 12.20 |
| Depression only (facilitation re-moved) | 10.58 ± 0.34 | 135.74 ± 5.37 | +0.010 ± 0.448 | −4.67 ± 6.08 |

Fits require finite, nonconstant activity, peak rate ≥ 0.2, *R*^2^ ≥ 0.5, width≥ 20 ms, and a complete 10–90% transition inside the window. Baseline and signed amplitude are bounded to [*−*1000, 1000], mid-point to [0, 600] ms, and *k* to [20*/*(2 log 9), 500] ms, with at most 200 function evaluations. Optimization exceptions and fits failing these criteria are excluded rather than assigned zero. Paired comparisons require the same threshold choice and the same neurons passing the criteria in both conditions. We average accepted paired cell-level changes within each trial, take the median across matched trials within each network, and report mean ± SEM across networks. Matched-trial counts per network, requiring at least one accepted cell pair, range from 1,205–1,700 for efficacy set to one, 877–1,841 for facilitation-only, and 1,277–1,671 for depression-only. All ten networks contribute to every sigmoid summary; no network is excluded based on output amplitude.

Removing depression increased the maximum absolute output in all ten networks, from 1.60–1.79 under intact STP to 1.92–1.71 *×*10^30^ under facilitation-only replay, with substantial variation in amplification across networks. Across all ten networks, normalized absolute slope increased in 8/10 and width decreased in 9/10. Some originally rising responses became falling or transient (Figure S4). Complete-trial output amplification is shown in Figure S5. The output amplification is consistent with a gain-limiting role of depression in these circuits and with synaptic gain control [Abbott et al., 1997].

Removing facilitation reduced the paired fitted amplitude in all ten networks (mean change 0.281 ± 0.088 rate units) and reduced the final target-minus-alternative output margin from 0.784 ± 0.013 to 0.319 ± 0.006, while normalized absolute slope changes were mixed (4/10 decreased).

##### Supplementary ramp and step fit scores

We retain the original linear-ramp and two-level-step fits and quantify their continuous *R*^2^ = 1 *−*SSE*/*SST scores. Row-wise rescaling prevents overflow and leaves *R*^2^ unchanged; constant or non-finite trajectories have no estimable score. Table 7 summarizes the full 0–1500-ms stimulus period; the original 0–600-ms window is retained as a window-sensitivity check. This analysis uses all recurrent neurons on common-correct threshold trials: even trial identifiers determine intact-network preference, and odd identifiers assess the same preferred-direction neuron/trial pairs with peak activity ≥0.2 in both conditions. Two facilitation-only networks lack preference-discovery trials for both directions and have no estimable paired scores; they are omitted from that score summary rather than assigned zero. Table 7 reports mean*±*SEM of within-network median paired score changes.

**Table 7:** Continuous fit-score changes over the full 0–1500-ms stimulus period. Values are mean ± SEM of within-network medians of paired neuron/trial *R*^2^ differences, using the original ramp and step fits. Network counts reflect availability under the original common-correct preference-discovery rule.

| Condition | Δ*R*<sup>2</sup><sub>ramp</sub> | Δ*R*<sup>2</sup><sub>step</sub> | Networks |
|---|---|---|---|
| Effective synaptic efficacy set to one | −0.063 ± 0.049 | −0.052 ± 0.018 | 10/10 |
| Facilitation only (depression removed) | −0.063 ± 0.020 | −0.011 ± 0.038 | 8/10 |
| Depression only (facilitation removed) | −0.148 ± 0.021 | −0.081 ± 0.013 | 10/10 |

**Table 8:**
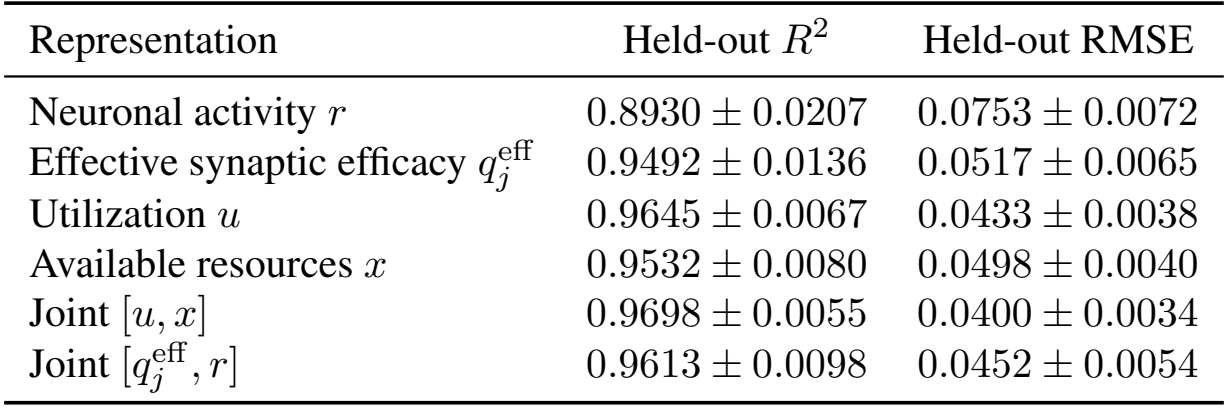
Shared-time decoding of cumulative input evidence. Values are mean*±*SD across ten trained networks. RMSE is in input units×s.

After facilitation removal, both ramp and step scores decreased in all ten networks over the full stimulus period, but only 8/10 and 5/10, respectively, in the primary 600-ms window. Depression removal did not consistently reduce both scores in either window.

##### Choice-output trajectories

Figure S5 compares both components on the same 2,000 nonzero-coherence trials per network, retaining all trials regardless of correctness or threshold crossing. Outputs are averaged within each network and stimulus direction, then summarized with equal network weights.

**Figure S4:**
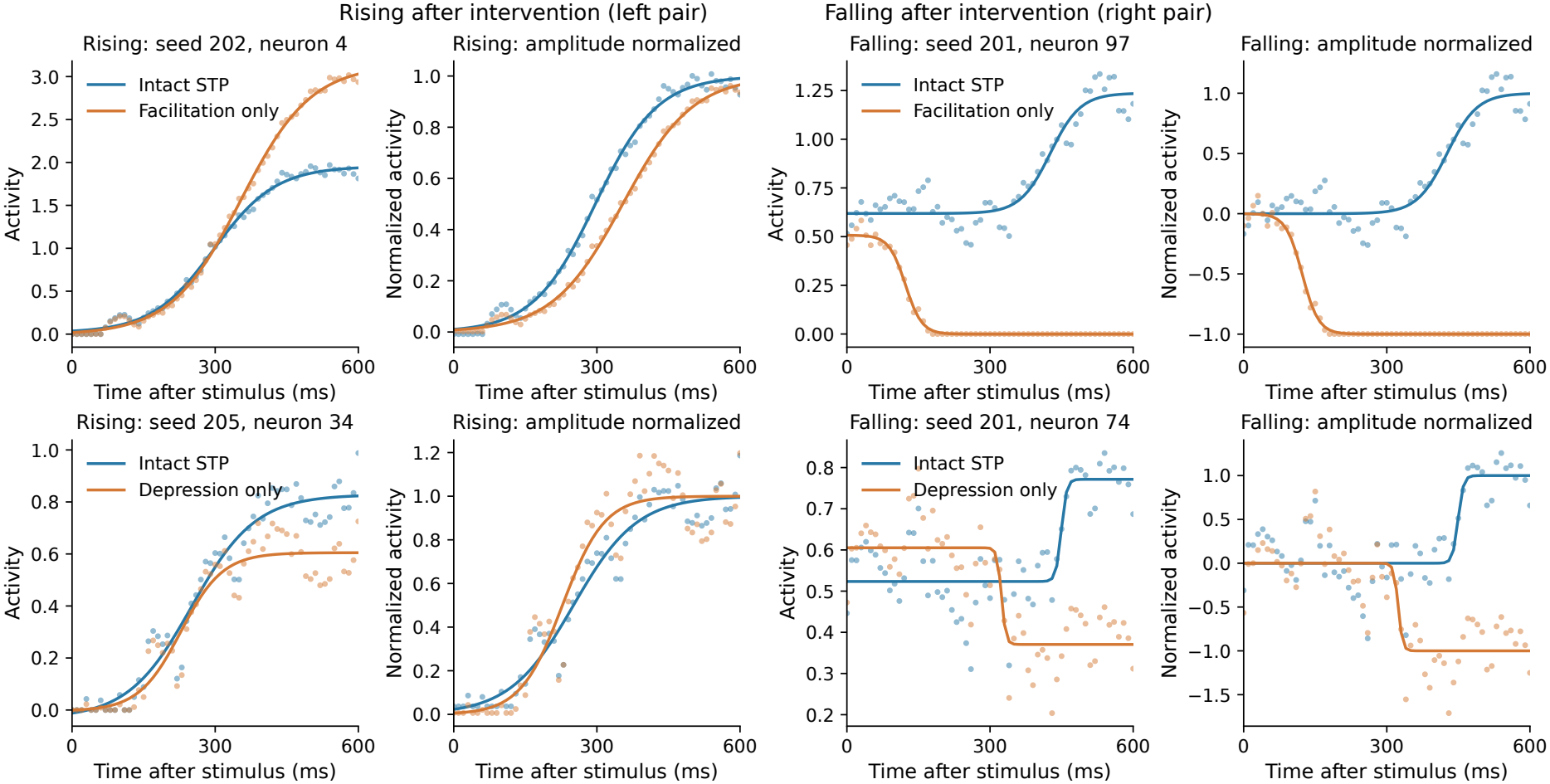
Paired neuronal responses grouped by direction after intervention. Top row: facilitation-only; bottom row: depression-only. Columns 1–2 show rising intervention responses and columns 3–4 show falling intervention responses. Each example overlays intact STP and the corresponding intervention on the same input and neuron, using matched threshold choices. Within each pair, the first panel shows raw activity and the second subtracts fitted baseline and divides by absolute fitted amplitude, preserving direction. Dots show activity and lines show accepted sigmoid fits during the first 600 ms after stimulus onset. Examples are stratified by fitted intervention-amplitude sign. Previously selected median-proximity examples are reused where available; added rising and falling examples retain the original fit criteria and are selected by paired fit quality within the first eligible batch or a previously recorded matching trial.

**Figure S5:**
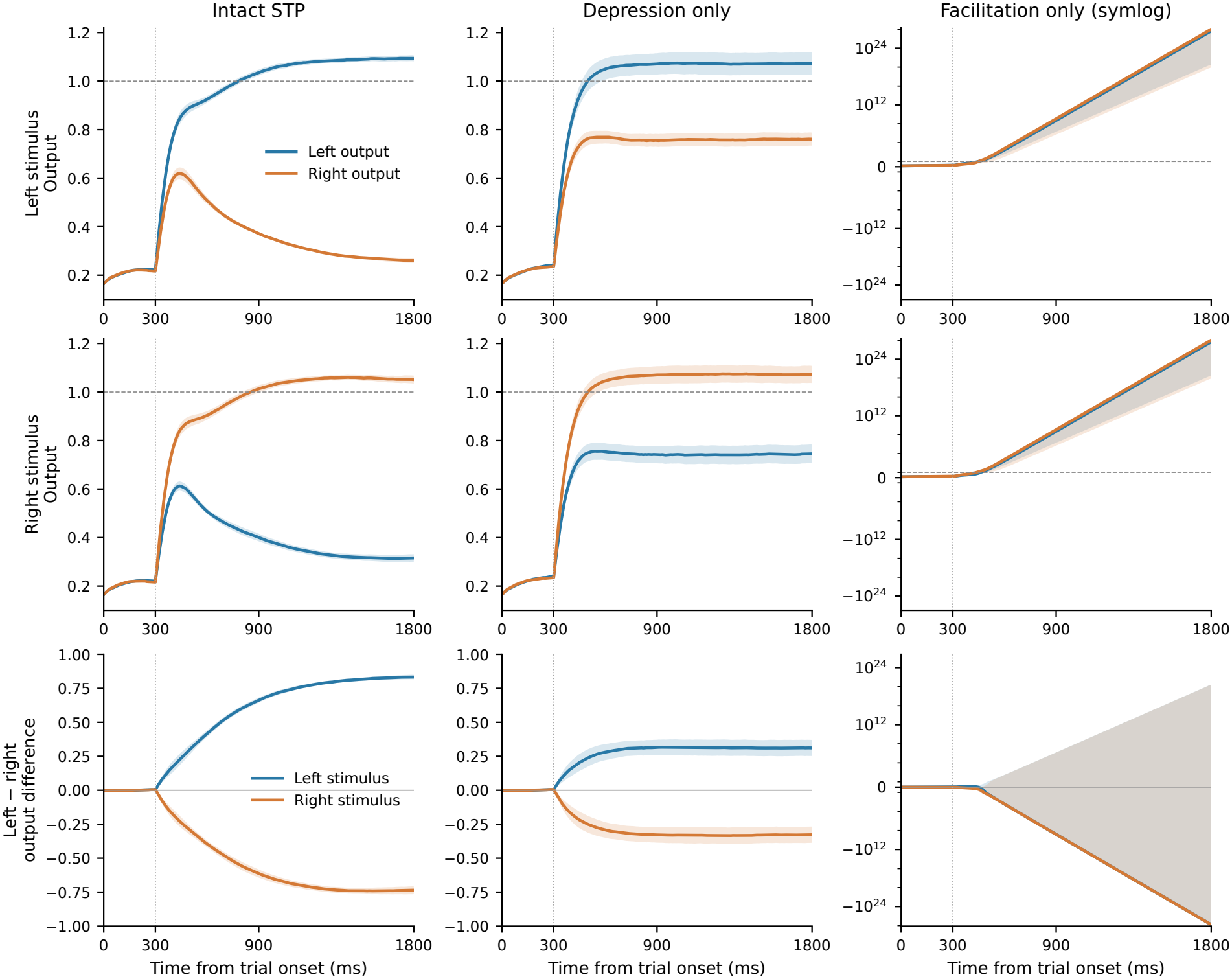
Output trajectories under component interventions, mean ± SEM across all ten networks. Columns show intact STP, depression-only, and facilitation-only replay. The first two rows show left and right output channels for left- and right-directed stimuli, respectively. The bottom row shows left-minus-right output differences for both stimulus directions; uncertainty is computed from within-network differences. Dotted lines mark stimulus onset at 300 ms and dashed lines mark the decision threshold of one. Intact and depression-only panels use linear axes; facilitation-only panels use a symmetric-logarithmic scale, linear between *−*1 and 1, to retain extreme finite outputs. Depression-only reduces output separation, whereas facilitation-only amplifies outputs to markedly different degrees across networks.

#### A.8 Decoding accumulated sensory evidence

We decoded the realized cumulative left-minus-right sensory input, including input noise, from neuronal activity *r*, effective synaptic efficacy 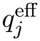 , utilization *u*, available resources *x*, and the concatenated states [*u, x*] and 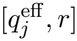 . Within each network, 1,600 complete trials formed the training set and 400 separate trials formed the held-out set. One ridge-linear decoder per representation (*λ* = 1) was fitted using all 75 sampled times, from 320 to 1,800 ms at 20-ms intervals; stimulus onset was 300 ms. The same decoder coefficients were applied at every held-out time, and all representations used the same trials and checkpoint within a network.

Both *u* and *x* outperformed *r* in all ten networks (mean paired Δ*R*^2^ = 0.0716 and 0.0603; two-sided Wilcoxon *p* = 0.00195 for each). Utilization exceeded resources in all ten (Δ*R*^2^ = 0.0113, *p* = 0.00195), and the joint [*u, x*] state gave the strongest readout. Effective efficacy 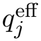 also exceeded activity in all ten networks and reduced RMSE by 31.2 ± 7.2% (mean ± SD of within-network reductions). Synaptic state therefore provided a more accessible linear readout of the population-level accumulated-evidence trend than neuronal activity.

#### A.9 Choice tuning and functional population assignment

Figure 3 uses ten RNNs with STP, all with mean initial STP strength 0.1 and training seeds 201–210. In these runs, an STP-strength learning-rate multiplier of 400 means that the raw parameter *θ*_*j*_ is updated at 400 times the base learning rate, while all other trainable parameters retain the base rate of 0.01. Each network is evaluated using its minimum-validation-task-loss checkpoint over its 100,000-update run. The multiseed analysis obtains the selected update from the checkpoint and applies that network’s tuning labels to all subsequent pruning and STP-strength analyses. These labels are estimated independently of training and of the perturbation evaluation stream. Each trained network is one independent replicate, and Figure 3 reports network means and SEM.

For each network, 300 left-evidence and 300 right-evidence trials are generated at 16% coherence, with 300 ms of fixation and 1500 ms of stimulus presentation. Opposite-direction trials are generated in pairs sharing the same input and recurrent noise realizations; the pair seeds are drawn using random seed 20260916. Simulation uses a 0.5-ms integration step and saves activity every 10 ms. A trial enters the tuning analysis if its first output threshold crossing (strictly above 1.0) indicates the correct direction. Rates are averaged over absolute trial time 1400–1800 ms, inclusive, giving 41 saved samples per trial. This is the final 400 ms of the trial, corresponding to 1100–1500 ms after motion onset while evidence remains present.

For each of the 100 neurons, the implementation applies a two-sided Welch test to the trial-level window means in the two conditions, with Benjamini–Hochberg correction across all neurons within that network. We denote the adjusted *p* value for neuron *j* by 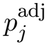 . Neurons with 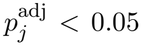 are classified as choice-selective; the sign of the left-minus-right mean defines preference. E/I identity then gives *E*_*L*_, *E*_*R*_, *I*_*L*_, and *I*_*R*_, and nonselective neurons remain in separate E and I groups. No effect-size or slope criterion enters this classification.

Figure 3b summarizes the same trials used for classification. Before cross-network aggregation, each network’s group means are divided by its maximum unit-level condition mean, without subtracting a minimum; empty groups are omitted. Across networks, 59.2 *±* 2.9% of excitatory neurons and 76.5 *±* 3.8% of inhibitory neurons were choice-selective (mean*±*SEM).

#### A.10 STP-strength distributions and matched neuronal silencing

Across networks, 98.6% of high-STP neurons were choice-selective, compared with 16.1% of low-STP neurons (Figure 3c). Accuracy was 85.7% in intact networks, 54.9% after high-STP neurons were silenced, and 82.0% after low-STP neurons were silenced (Figure 3d). Accuracy after low-STP silencing exceeded that after high-STP silencing by 27.1 percentage points on average (95% bootstrap confidence interval, 17.2–36.0; exact paired-randomization *p* = 0.004).

STP strength in Figure 3c denotes the learned *a*_*j*_. During evaluation, *a*_*j*_ remains constant, while normalized synaptic efficacy *u*_*j*_*x*_*j*_*/U* and the recurrent transmission factor 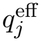 evolve within each trial. STP strengths are histogrammed separately for E and I neurons in bins of width 0.02. For display, each network’s bin counts are smoothed with a Gaussian kernel of standard deviation one bin and radius three bins, with edge-value padding, then summarized as network means and SEM. The ordinate reports neuronal counts per network. High-STP membership is *a*_*j*_ ≥ 0.5 and low-STP membership is *a*_*j*_ *<* 0.5. At the selected checkpoints, high-STP neurons accounted for 56.1 ± 10.5% of all neurons, 53.8 ± 10.0% of E neurons, and 65.5 *±* 15.4% of I neurons (mean *±* SD across networks). Choice-selective fractions use every member of the corresponding threshold-defined group in each network, and are then averaged over networks with equal weight. The choice-selective fractions were 98.7% versus 12.5% for high-versus low-STP E neurons and 98.1% versus 35.0% for the corresponding I groups. The high-versus-low contrast was insensitive to the exact STP-strength cutoff: with cutoffs of 0.4, 0.5, and 0.6, the across-network choice-selective fractions were respectively 98.6% versus 15.9%, 98.6% versus 16.1%, and 98.5% versus 17.1%. This sensitivity analysis changes only the grouping used in Figure 3c; the matched silencing subsets in Figure 3d retain the primary cutoff of 0.5 and their deterministic extreme-STP-strength selection.

Figure 3d uses matched subsets of those groups. Within each network and each cell type *c* ∈ {*E, I }*, let *n*_*c*_ = min(*N*_*c*,high_, *N*_*c*,low_). The silencing comparison selects the *n*_*c*_ largest STP strengths from the high group and the *n*_*c*_ smallest STP strengths from the low group. Combining the E and I subsets matches both total count and E/I composition. Selection is deterministic rather than random sampling; the individual panel reports a rounded mean of 40 silenced neurons per network in each condition.

For a selected set *S*, silencing is imposed throughout the trial as

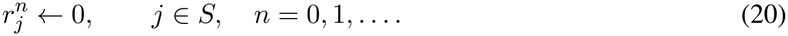

This assignment is applied at initialization and after every neural-state update. Recurrent and readout contributions from selected neurons are therefore removed, while the neural state equations and STP updates continue to run. The analysis evaluates intact networks and networks with the matched high- or low-STP subsets silenced, using the same behavioral stream and accuracy definition as pruning. The paired-randomization test uses the within-network accuracy difference between low-STP and high-STP silencing conditions and follows the procedure specified in Appendix A.3.

#### A.11 Connection selectivity and STP strength

For each network, we correlated learned STP strength with three weight measures using Spearman’s *ρ*. Sensory-input selectivity is 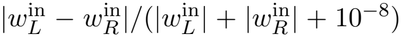 choice-output selectivity uses the corresponding two readout weights. Outgoing recurrent-weight magnitude is the *L*_2_ norm of a source neuron’s column in *W* ^rec^. Table 9 gives the mean within-network correlation; two-sided Wilcoxon tests use the ten network correlations. Figure 3e visualizes the association between STP strength and sensory-input selectivity. For this visualization, sensory-input selectivity is averaged within the low- and high-STP groups in each network, using the strength cutoff of 0.5 defined in Appendix A.10. The Spearman correlations use all neurons within each network before the network-level coefficients are summarized.

**Table 9:** Trained connectivity and STP strength across ten networks. Selectivities are unsigned normalized left–right differences.

| Weight measure | Mean Spearman ρ | Consistent networks | Two-sided *p* |
|---|---|---|---|
| Sensory-input selectivity | +0.574 | 10/10 positive | 0.00195 |
| Outgoing recurrent-weight *L*<sub>2</sub> norm | −0.149 | 10/10 negative | 0.00195 |
| Choice-output selectivity | +0.064 | 7/10 positive | 0.160 |

Stronger STP is associated with more selective sensory input, whereas its association with recurrent-weight magnitude is negative and its association with readout selectivity is weak.

#### A.12 Sensory, readout, and recurrent connection pruning

The pruning experiments hold the checkpoint and functional group assignments fixed and set the specified weight block to zero during forward evaluation, without retraining. Within a network, all variants share the same inputs and recurrent noise. Each evaluation uses random seed 100, with 1500 trials for each of the ten nonzero coherence–direction conditions (*c* = 1, 2, 4, 8, 16% in both directions), plus 1500 zero-coherence trials. Thus 16,500 trials are simulated per variant; overall accuracy is the fraction of correct final-output argmax choices among the 15,000 nonzero-coherence trials. Threshold timeouts are retained in this denominator and judged by the final output. Simulation uses a 0.5-ms step and 10-ms output sampling.

Let 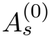 and 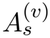 be intact and pruned accuracies, expressed as fractions, in training seed *s*. Heatmaps

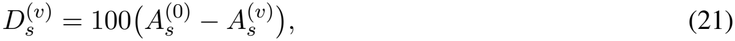

in percentage points. Positive values indicate impaired performance and negative values indicate improvement.

The three connection-pruning analyses are:

1. **Sensory input (Figure 3f)**. Set 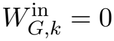 for each of the six populations *G* and each sensory channel *k* ∈ {0, 1 } , giving 12 pruned conditions plus the intact condition. The cue channel remains connected. Because the complete input weight is removed, this operation deletes both the coherence-dependent signal and the noisy baseline-offset contribution carried by that channel.
2. **Choice readout (Figure 3g)**. Disconnect each of the three E populations from both outputs in turn by setting 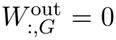 direct I-to-output weights are structurally absent. Effects are evaluated separately on 7500 left-evidence and 7500 right-evidence trials and referenced to intact accuracy on the same evidence side. The intervention removes the connections from the source population to both outputs.
3. **Recurrent pathways (Figure 3h)**. For every ordered pair of the four tuned populations, remove the connections from source *G* to target *H* by setting the complete block *W*_*H,G*_ = 0, giving 16 pruned conditions plus intact. Within-group blocks are included, nonselective neurons remain active, and blocks involving nonselective groups are not tested. Empty blocks produce a recorded effect of zero.

Each 2 *×* 2 E/I matrix in Figure 3h uses an independent color scale and displays the mean ± SEM effect magnitude for each connection block. Outlines mark the largest across-network mean loss among the three off-diagonal targets for each source and, separately, the two largest mean effects in the E-to-E submatrix.

### B Implementation and stability analysis of the theoretical model

The simulations use the four-population model defined in Equations 5–8. The model describes population firing rates and effective population-to-population synaptic efficacy, inspired by the trained RNN’s choice-selective E–I organization and recruitment of STP.

#### B.1 Simulation and parameterization

For cell class *a* ∈ *{E, I}*, the activation function is

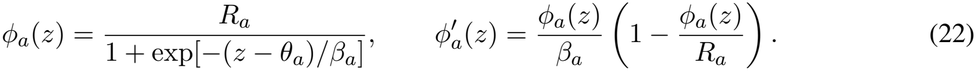

Firing rates are in Hz and time is in seconds. Inputs use the model’s dimensionless input scale, so *J*_*EE*_, *J*_*EI*_, and *J*_*IE*_ convert rates into that scale. Parameter values are listed in Table 10.

**Table 10:**
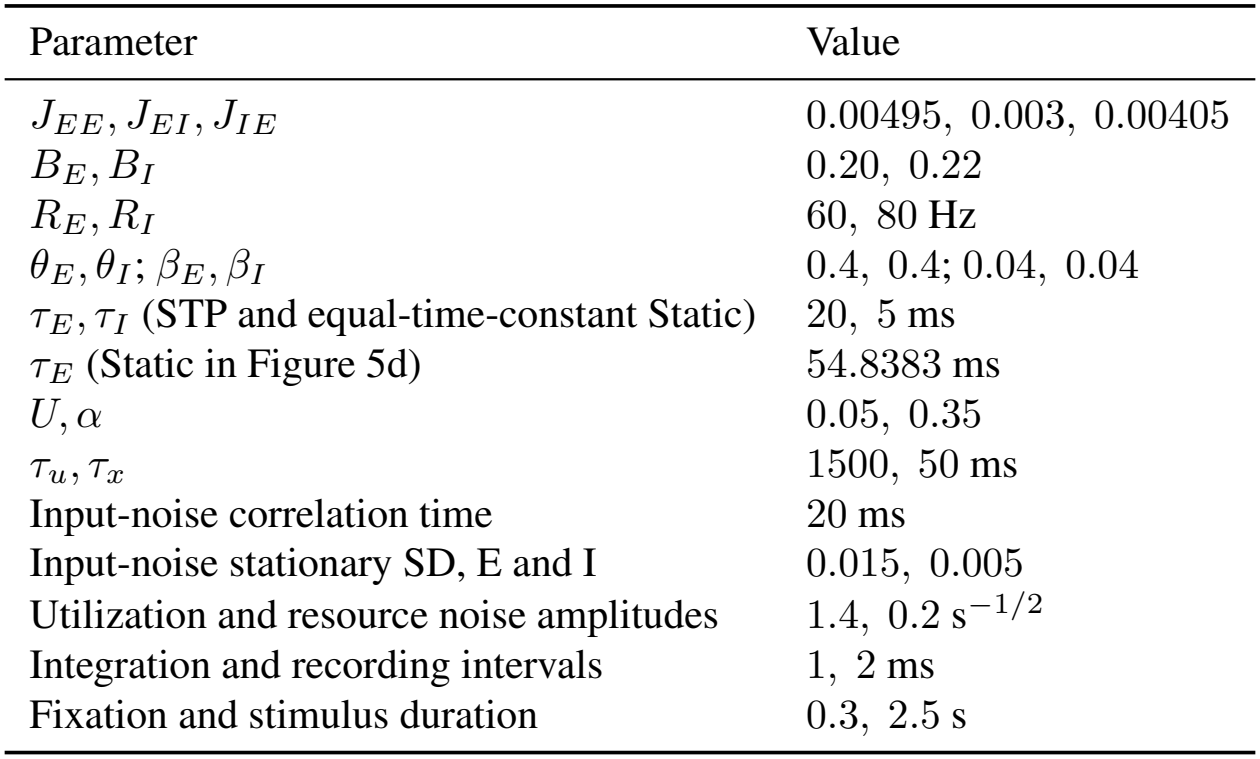
Parameters of the four-population theoretical model.

**Table 11:**
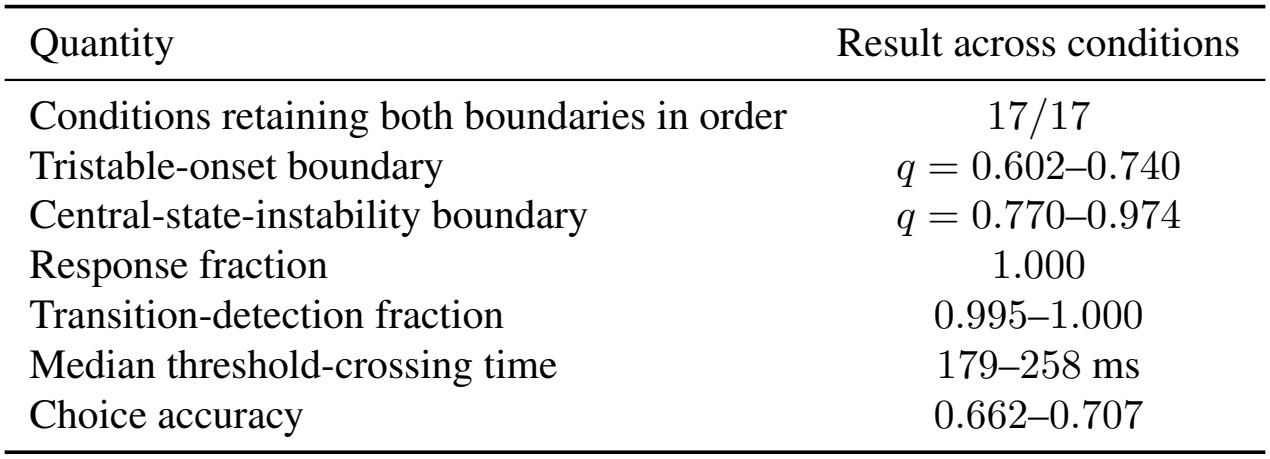
Finite-grid robustness of the four-population mechanism. Ranges are taken across the 17 prespecified conditions; no condition was retuned. Accuracy counts unresolved responses as incorrect.

**Table 12:** Behavior-matched activity cost of the theoretical STP and Static circuits. Paired entries are STP/Static. Reductions are relative to Static; 95% confidence intervals use 20,000 paired bootstrap resamples of independent 200-trial noise batches.

| Target tier | Accuracy | Mean RT (ms) | Expected spikes | Reduction (95% CI) |
|---|---|---|---|---|
| 85% | 84.24/83.80% | 264.9/266.6 | 4.723/6.567 | 28.08% (27.72–28.46) |
| 90% | 89.01/88.70% | 410.2/414.8 | 7.675/10.269 | 25.26% (24.84–25.70) |
| 95% | 94.45/94.83% | 747.4/748.9 | 15.160/18.472 | 17.93% (17.69–18.17) |

During the stimulus, *e*_*L*_ = 0.1+0.01*c* and *e*_*R*_ = 0.1 0.01*c*, where positive *c* favors left. Evidence inputs are zero during fixation. Each population receives Ornstein–Uhlenbeck input noise with the correlation time and stationary SD in Table 10. The excitatory noise terms are independent; the two inhibitory populations receive the same external noise realization in the displayed simulations. Their neuronal activities remain separate variables.

The deterministic drift is integrated by the fourth-order Runge–Kutta method. After each step, independent Gaussian increments are added to the log odds of each excitatory *u*_*p*_ and *x*_*p*_, with SD proportional to the square root of the integration interval; this corresponds to bounded Stratonovich noise for *z* ∈ {*u*_*p*_, *x*_*p}*_ . Inhibitory synapses have no STP or synaptic noise. Each simulation starts from the no-stimulus static neuronal equilibrium with *u*_*p*_ = *U* and *x*_*p*_ = 1, followed by the fixation period. Static simulations use *q*_*L*_ = *q*_*R*_ = 1, while the STP model starts at *q*_*L*_ = *q*_*R*_ = 0.35. The robustness claim in Section 5.1 was verified by rerunning the timing and decision35 protocols with STP enabled on all four populations’ outputs and applying *q* to all four populations in the frozen-synapse scan: the tristable-onset and center-instability boundaries are unchanged (0.664 and 0.862), and behavior is comparable (accuracy 65.0% versus 66.2%, median threshold-crossing time 197 ms versus 233 ms, mean spike count 4.24 versus 4.14).

#### B.2 Transition detection and stability analysis

A descriptive transition time in Figure 4b–d is fitted to the winning excitatory trace using two constant activity levels. Both intervals must last at least 100 ms, the fitted increase must be at least 15 Hz, and the explained variance must be at least 0.5. Figure 5c–e instead uses the first crossing of |*r*_*E,L*_ *− r*_*E,R*_| = 35 Hz. These are distinct alignment and response conventions. In the original model, the winner’s synaptic gain crosses the center-instability boundary before the neuronal transition: over the 200 trials of the Figure 4 protocol, the winning population’s *q*_*p*_ = *αu*_*p*_*x*_*p*_*/U* crosses 0.862 at a median of 137 ms, whereas the fitted activity transition occurs at a median of 228 ms, and the losing population’s gain stays near the tristable onset (Figure S7).

The synaptic states also retain signed input history during the low-activity period. A linear decoder on the utilization and resource differences (*u*_*L*_ *− u*_*R*_, *x*_*L*_ *− x*_*R*_) recovers evidence direction above chance throughout the silent period, rising from 57.6% at 100 ms after stimulus onset to 64.4% at 400 ms (five-fold cross-validated linear discriminant analysis; 1600 trials over eight nonzero coherence levels, 200 trials per level and five seeds; chance 50%).

Psychometric curves (Figure S6), which measure choice accuracy as a function of evidence strength, use seeds 100–104 with 40 trials per coherence level, giving 2200 trials per condition. The total-rate comparison of Figure 5d uses an adjusted variant of the STP model in which the noise added to the synaptic states is set to zero, with all other parameters unchanged. With synaptic-state noise, the STP model has lower psychometric accuracy than the Static model at these operating points. Removing this noise yields overlapping 95% Wilson intervals for the STP and Static psychometric curves at every evidence level.

For stability analysis, we remove noise, apply equal stimulus inputs, and hold *q*_*L*_ = *q*_*R*_ = *q* fixed. With the ordering (*r*_*E,L*_, *r*_*E,R*_, *r*_*I,L*_, *r*_*I,R*_), define

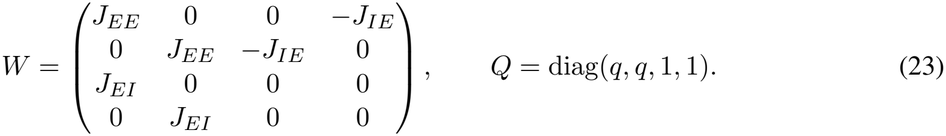

At an equilibrium **r**^*^, let *D*_*ϕ*_ be the diagonal matrix of activation-function derivatives at the four population inputs and *D*_*τ*_ = diag(*τ*_*E*_, *τ*_*E*_, *τ*_*I*_, *τ*_*I*_). The neuronal Jacobian is

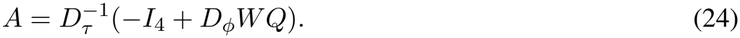

An equilibrium is stable if all four eigenvalues have negative real parts. The recorded equilibrium search spans *q* = 0.35–1.5 in increments of 0.002, combining scalar bracketing for symmetric roots with multiple initial guesses for asymmetric roots. The maximum recorded equilibrium residual is below 3.8 *×* 10^*−*12^. At *q* = 0.35, the stable symmetric equilibrium has excitatory rates of approximately 5.1 Hz. The sampled data first show three stable equilibria at *q* = 0.664 and loss of stability of the symmetric equilibrium at *q* = 0.862. These values locate the regimes at the resolution of the parameter sweep.

At a symmetric equilibrium, write 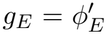 and 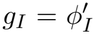 . The differences *d*_*E*_ = (*r*_*E,L*_ *− r*_*E,R*_)*/*2 and *d*_*I*_ = (*r*_*I,L*_ *− r*_*I,R*_)*/*2 obey the linearized equation

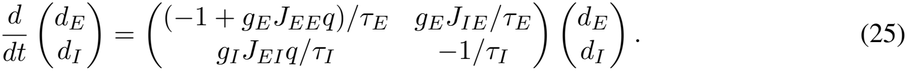

The positive off-diagonal terms express opponent inhibition: an increase in *I*_*L*_ relative to *I*_*R*_ favors *E*_*L*_ by suppressing *E*_*R*_. A zero eigenvalue in this mode satisfies *g*_*E*_*q*(*J*_*EE*_ + *g*_*I*_*J*_*IE*_*J*_*EI*_) = 1. Both activation-function derivatives depend on the equilibrium. The stability calculation evaluates the common-activity mode and the other equilibria in addition to this choice mode.

**Figure S6:**
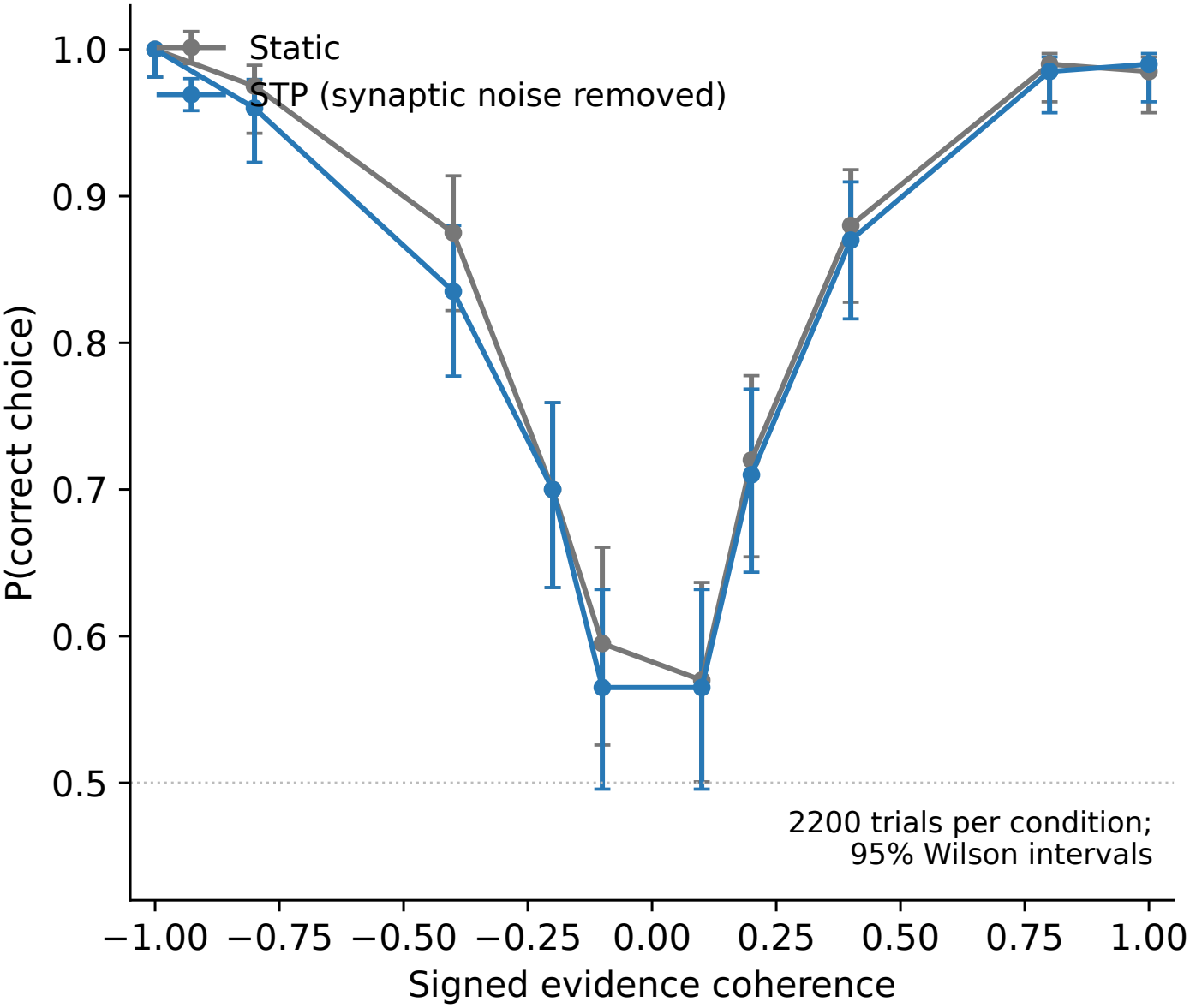
Psychometric comparison of the Static model and the adjusted STP variant used in Figure 5d (parameters adjusted for a matched psychometric curve; all other parameters unchanged). Fraction of correct choices as a function of signed evidence coherence; error bars show 95% Wilson intervals over 2200 trials per condition (11 coherence levels, 40 trials per level, five seeds). The curves overlap at every evidence level, characterizing the accuracy of the operating points displayed in Figure 5d. The controlled cost comparison in Figure 5e is specified separately in Appendix B.5.

**Figure S7:**
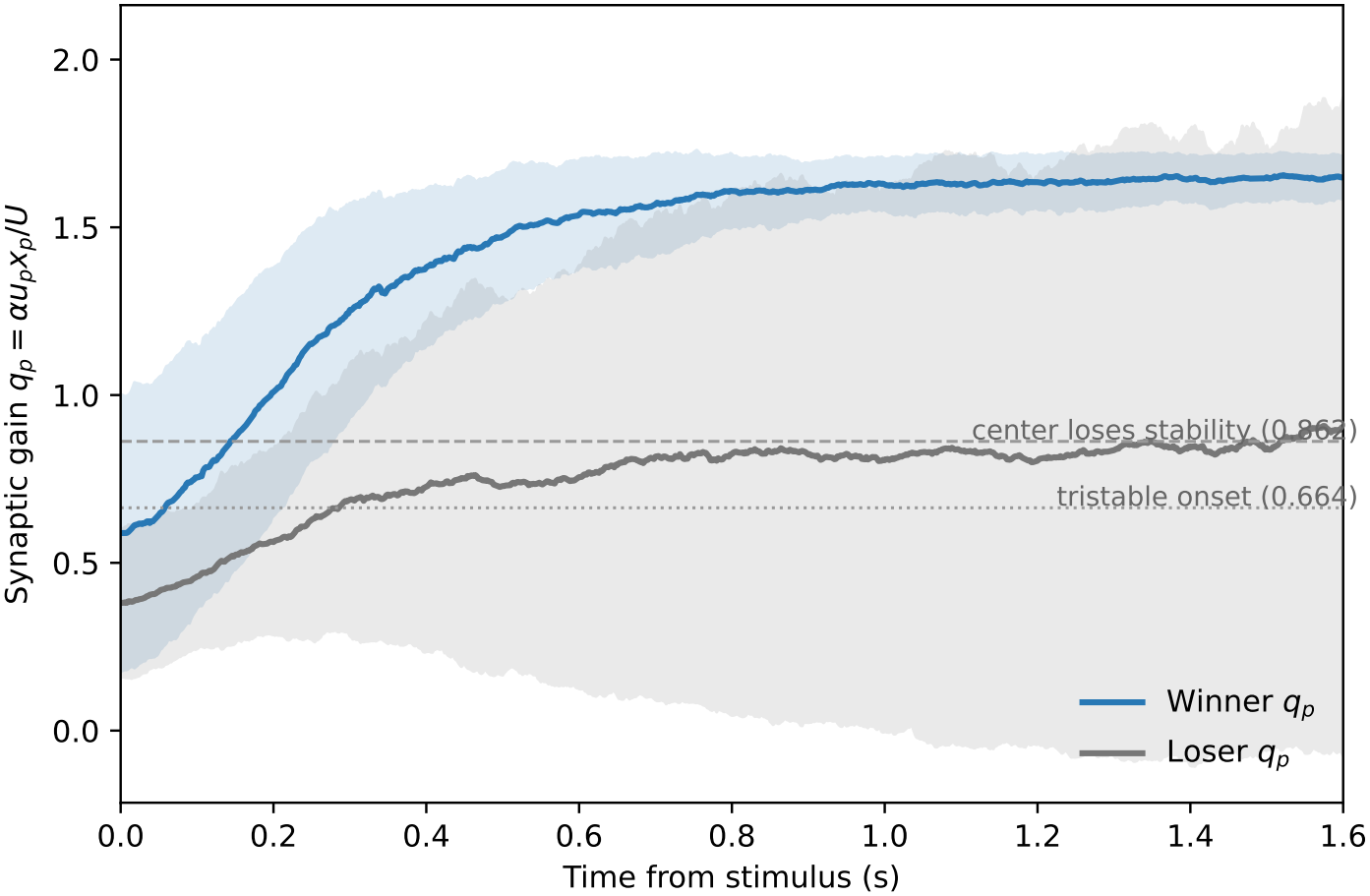
Synaptic gain relative to the frozen-synapse stability boundaries. Mean ± SD of *q*_*p*_ = *αu*_*p*_*x*_*p*_*/U* for the winning and losing populations across the 200 trials of the Figure 4 protocol in the original model. The winner’s gain crosses the center-instability boundary (0.862) at a median of 137 ms, before the median activity transition (228 ms); the loser’s gain stays near the tristable onset (0.664).

#### B.3 Parameter-region robustness and slow–fast interpretation

The parameters define a population circuit in which the effective synaptic gain can move the faster neuronal subsystem from a stable low-activity regime to a choice-selective regime. The baseline frozen-gain scan samples *q* = 0.35–1.5 in increments of 0.002. To test secondary-parameter sensitivity without condition-specific retuning, we additionally evaluated a prespecified 17-condition grid: a 3 *×* 3 grid of *−*10%, baseline, and +10% recurrent self-excitation and inhibitory-to-excitatory coupling (nine conditions, including the baseline); two ±10% changes in excitatory-to-inhibitory coupling; and 0.5-fold or 2-fold changes in each of *U* , *τ*_*u*_, and *τ*_*x*_. Each dynamic condition used 1000 trials across five noise seeds.

The preserved boundary order and transition detection show that the mechanism occupies a finite region of circuit and STP parameters.

The frozen-synapse portrait is interpreted through a quasi-steady-state, slow–fast approximation, a standard analysis for systems with neuronal and synaptic variables evolving on different time scales [Hennig, 2013, Tsodyks and Wu, 2013]. Let *τ*_*n*_ denote the neuronal time scale and define *ϵ*_*u*_ = *τ*_*n*_*/τ*_*u*_ and *ϵ*_*x*_ = *τ*_*n*_*/τ*_*x*_. On the fast time *s* = *t/τ*_*n*_, the coupled system can be written schematically as

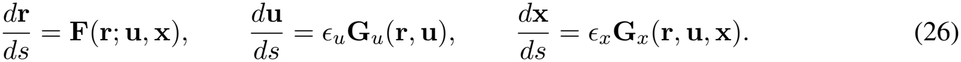

For each fixed (*u*_*p*_, *x*_*p*_) state, setting **F** = 0 defines the instantaneous equilibrium manifold of the faster neuronal subsystem, whose local stability is given by Equation 24. The evolving synaptic state then moves the neuronal subsystem through this family of phase portraits. The frozen portrait identifies the stability boundary approached by the dynamic trajectory, and the dynamic simulation shows that the winner’s effective gain crosses this boundary before the fitted neuronal transition (Figure S7).

#### B.4 Expected-spike-count comparison

For Figure 5d,e, we define the total firing rate and expected spike count as

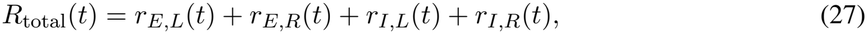

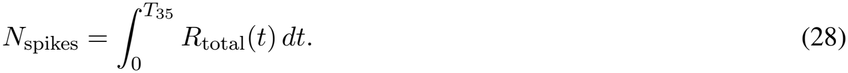

Here, *T*_35_ is the first crossing of |*r*_*E,L*_ *− r*_*E,R*_| = 35 Hz. All four rates contribute positively, including activity in the losing choice populations. Figure 5d displays threshold-aligned total rates at psychometrically matched operating points; Figure 5e compares expected decision-period spikes at three accuracy- and reaction-time-matched tiers (Appendix B.5). The integration uses linear interpolation at the crossing time. The original total-rate comparison in Figure 5d uses five batches of 200 trials per condition with independent noise seeds. The three-tier comparison in Figure 5e uses the separately calibrated models and trial counts specified in Appendix B.5.

Each population is represented by one effective rate unit in this calculation. Because rates are in Hz and time is in seconds, their integral gives the expected spike count of the four-unit model. For population index *a* ∈ *{E*_*L*_, *E*_*R*_, *I*_*L*_, *I*_*R*_*}*, let *n*_*a*_ denote population size and *r*_*a*_ its mean firing rate. Then the corresponding network spike count would be ∫∑_*a*_ *n*_*a*_*r*_*a*_(*t*) *dt*. Both conditions use the same unit population weights.

The Static value is *τ*_*E*_ = 54.8383 ms, the STP value is *τ*_*E*_ = 20 ms, and both models use *τ*_*I*_ = 5 ms. The Static time constant was calibrated using 1000 separate trials (noise seed 510) to match the rise time of the stimulus-aligned mean excitatory activity difference of the original STP model from 12 to 48 Hz; these levels correspond to 20% and 80% of a fixed 60-Hz reference. The calibrated rise durations are approximately 226 ms for Static and 221 ms for STP in the original model. In the adjusted variant used for Figure 5d, the STP rise duration is approximately 145 ms, compared with 226 ms for Static; these operating points have overlapping psychometric curves (Figure S6). Evaluation uses seeds 410–414 for both the original model and the adjusted variant. All evaluation trials in the original displayed operating points reach the decision threshold.

#### B.5 Behavior-matched comparison of decision-period activity cost

We compared the four-population STP and Static models at three target-accuracy tiers (85%, 90%, and 95%). Stimulus coherence had fixed magnitude 0.4 with balanced left and right directions, and paired trials shared the stimulus and input noise. Excitatory neuronal time constants were calibrated separately for the two models and then frozen before evaluation. The 85% and 90% tiers were selected by accuracy; the 95% tier jointly matched accuracy and mean reaction time. We evaluated 8,000 independent paired trials per model at each of the first two tiers and 16,000 at the third. All evaluation trials reached the 35-Hz choice threshold; correct and incorrect choices both contribute to the activity measure.

Expected spikes are the integral of the sum of the four population rates from stimulus onset to that model’s own threshold crossing, with one effective rate unit per population (Appendix B.4). Figure 5e and Table 12 report the held-out results. The excitatory time constants for STP/Static were 14.107/41.604 ms, 39.692/64.101 ms, and 123.610/114.877 ms at the three tiers. Thus, this comparison matches behavior while allowing internal time constants to differ.

Observed accuracy differed by at most 0.44 percentage points and mean RT by 0.19–1.10%. Paired 90% confidence intervals for these differences were contained within equivalence ranges of ±2 percentage points and ±15 ms. STP required fewer expected spikes at every tier, including the highest tier where mean RT differed by only 1.46 ms. This supports a lower spike-related activity cost at comparable decision performance.

## Notes

### Competing Interest Statement

The authors have declared no competing interest.

### Summary of Updates

This version expands the analysis of short-term synaptic plasticity (STP) in the trained recurrent networks. The Results and Methods now include fixed-weight STP interventions, decoding of accumulated sensory evidence, parameter-sensitivity analyses, and additional tests of the relationship between STP strength, choice selectivity, and network connectivity. Figures 2 and 3 have been revised to present these analyses. Figure 5 now includes a behavior-matched activity-cost comparison and expanded stability and robustness results. The Appendix and supplementary materials have also been updated with additional methodological details, component-ablation analyses, neuronal-transition examples, output trajectories, and revised references.

